# Microbial diversity in tropical marine sediments assessed using culture-dependent and culture-independent techniques

**DOI:** 10.1101/2021.02.27.433211

**Authors:** Alyssa M. Demko, Nastassia V. Patin, Paul R. Jensen

## Abstract

The microbial communities associated with marine sediments are critical for ecosystem function yet remain poorly characterized. While culture-independent (CI) approaches capture the broadest perspective on community composition, culture-dependent (CD) methods can capture low abundance taxa that are missed using CI approaches. The aim of this study was to assess microbial diversity in tropical marine sediments collected from five shallow water sites in Belize using both CD and CI approaches. CD methods captured approximately 3% of the >800 genera detected across the five sites. Additionally, 39 genera were only detected using CD approaches revealing rare taxa that were missed with the CI approach. Significantly different communities were detected across sites, with rare taxa playing an important role in the delineation of sediment communities. This study provides important baseline data describing shallow water sediment microbial communities and evidence that standard cultivation techniques may be more effective than previously recognized.

**Originality-Significance Statement:** Marine sediments host some of the most diverse microbial communities on the planet. While these communities are critical for global nutrient cycling, the oceanic food web, and the maintenance of ecosystem dynamics, they remain poorly studied. Studies that have assessed sediment communities typically use culture-independent approaches, which have known biases and can miss ecologically important taxa. Here we describe microbial diversity in marine sediments using both culture-dependent and culture-independent approaches. Our culturing approach, sequencing communities as opposed to individual colonies, revealed an additional 39 genera that were not detected with culture-independent methods. Additionally, we cultured numerous, as-yet undescribed species, suggesting that traditional culturing practices can be more efficient than commonly thought. Moreover, our results indicate rare taxa play an important role in distinguishing microbial communities at different sites, thus highlighting the importance of deep sequencing and incorporating culture-dependent approaches for diversity assessments.

## Introduction

Advances in sequencing technologies and bioinformatics have led to major improvements in our ability to assess the diversity and distributions of environmental microbes (bacteria and archaea) (Lynch and Neufeld, 2015; Hug *et al*., 2016; Thompson *et al*., 2017). The application of culture-independent (CI) methods has transformed our understanding of microbial diversity while metagenome assembled genomes and single cell genomics have provided insight relevant to functional traits in yet to be cultured organisms (Kalisky and Quake, 2011; Evans *et al*., 2015; Parks *et al*., 2017). Despite these advances, the microbial diversity associated with marine sediments remains poorly characterized relative to other major biomes such as soil and seawater (Lloyd *et al*., 2018; Martiny, 2019; Baker *et al*., 2021). Sediment microbial communities are diverse (Thompson *et al*., 2017), densely populated, (Dale, 1974; Musat *et al*., 2006), play integral roles in fundamental ecosystem processes (Snelgrove *et al*., 1997; Baker *et al*., 2021), and can exhibit extraordinary levels of fine-scale spatial structure (Probandt *et al*., 2018). While the inaccessibility of deep-sea sediments (>200 m depth) may contribute to the lack of data, the communities associated with shallow euphotic (≤200 m depth) water sediments also remain poorly described (Baker *et al*., 2021). Given that marine sediments cover ∼70% of the earth’s surface (Parks and Sass, 2009), baseline information describing sediment microbial diversity provides an important mechanism to understand community structure over time and across environmental gradients.

The gains afforded by CI diversity estimates can overshadow the intrinsic value of microbial cultivation. Culture-dependent (CD) methods provide opportunities to assess microbial metabolism and contributions to ecosystem function in ways that cannot be achieved using CI approaches. For instance, cultivation of *Nitrospira* provided critical insight into the first bacterium known to perform complete nitrification (Daims *et al*., 2015) while culture-dependent research with *Thermosulfidibacter takaii* ABI70S6^T^ resulted in the discovery of a reversible TCA cycle that was not detected with metagenomics (Nunoura *et al*., 2018). Additionally, testing for inhibition among marine *Vibrio* strains revealed that competition is greater between than within ecologically cohesive populations (Cordero *et al*., 2012) while CD work on two closely related species of *Salinispora* demonstrated ecological trade-offs in competitive strategies (Patin *et al*., 2015). Culturing techniques have been developed based on metabolic requirements inferred using CI techniques (Tripp *et al*., 2008). Thus, these approaches can offer complementary insights into microbial ecology, with the general observation that CI techniques provide more comprehensive taxonomic coverage while CD methods can provide clearer taxonomic resolution (Orphan *et al*., 2000; Chen *et al*., 2008; Shivaji *et al*., 2011; Vaz-Moreira *et al*., 2011; Dickson *et al*., 2014). While both techniques have been used in tandem, surprisingly few studies have focused on major environmental biomes such as marine sediments. Of note, the importance of media specificity and rare taxa when comparing CI and CD approaches was recently reported (Pédron *et al*., 2020).

Recent bioinformatic comparisons of CD versus CI bacterial diversity have revived discussion of the “great plate count anomaly” and the canonical theory that fewer than one percent of bacterial taxa have been cultured (Martiny, 2019; Steen *et al*., 2019). These studies provide contrasting views on the improvements that have been made in culturing relative to the proportions of bacteria that remain uncultured (Martiny, 2019; Steen *et al*., 2019). Advances such as these highlight the need to reassess frequently cited paradigms describing bacterial culturability, including the great plate count anomaly. While the majority of bacterial taxa have yet to be cultured (Lloyd *et al*., 2018), the development of innovative techniques in combination with persistent use of traditional methods has led to the successful cultivation of notable microbes including bacterioplankton in the SAR11 clade (Rappé *et al*., 2002; Henson *et al*., 2018), the first Asgard archaea representative ‘*Candidatus* Prometheoarchaeum syntrophicum’ (Imachi *et al*., 2020) and three Saccharibacteria (TM7) species with their Actinobacteria host (Cross *et al*., 2019), suggesting that many if not most microbes can ultimately be brought into the laboratory (Lewis *et al*., 2020).

The aim of this study was to assess microbial diversity in marine sediments using both CD and CI techniques. CD sample communities were determined by sequencing environmental plates inoculated from sediments while CI sample communities were determined by direct sequencing of sediments. Through the use of next-generation sequencing, we determined that culturing efficiency was 1-2% based on the number of 16S rRNA amplicon sequence variants (ASVs) detected and 3-4% in terms of the number of genera detected. Additionally, the CD method detected 39 genera that were not detected using the CI approach, highlighting the importance of culturing for capturing rare members of the community. These results emphasize the importance of both CD and CI methods for assessments of microbial diversity in marine sediments.

## Results

### Culture-Dependent (CD) Sediment Microbial Diversity

Across all five sites, CD alpha diversity analyses revealed on average 54 ± 3 16S rRNA amplicon sequence variants (ASVs) for sediments plated on seawater agar (SWA) and 39 ± 2 ASVs for the same sediment samples plated on marine agar (MA) (Figure S1a). SWA yielded significantly higher phylogenetic richness (Figure S2a; Faith’s PD Kruskal-Wallis H=8.870, p=0.003) and greater evenness (Figure S2b; Pielou’s Kruskal-Wallis H=7.645, p=0.006) when compared to the ASVs detected on MA.

The microorganisms detected on the SWA and MA plates were classified into five bacterial phyla (Figure 1a, Figure S3a). The three most abundant phyla were similar for both media, with Proteobacteria being the most abundant, averaging 80% and 96% of the communities on SWA and MA plates, respectively. Bacteroidetes represented 20% of the SWA communities compared to ca. 1.6% for MA, indicating that the low nutrient medium was more selective for this phylum. On MA, Firmicutes was the second most abundant phylum at ca. 2.6%. In comparison, Firmicutes averaged ca. 0.13% on SWA, suggesting that the nutrient rich MA media better selects for this phylum. The other two phyla detected in culture were Epsilonbacteraeota (formerly Epsilonproteobacteria) and Actinobacteria, both of which averaged <1% of the community in SWA and MA (Figure 1, Figure S3a).

**Figure 1.**
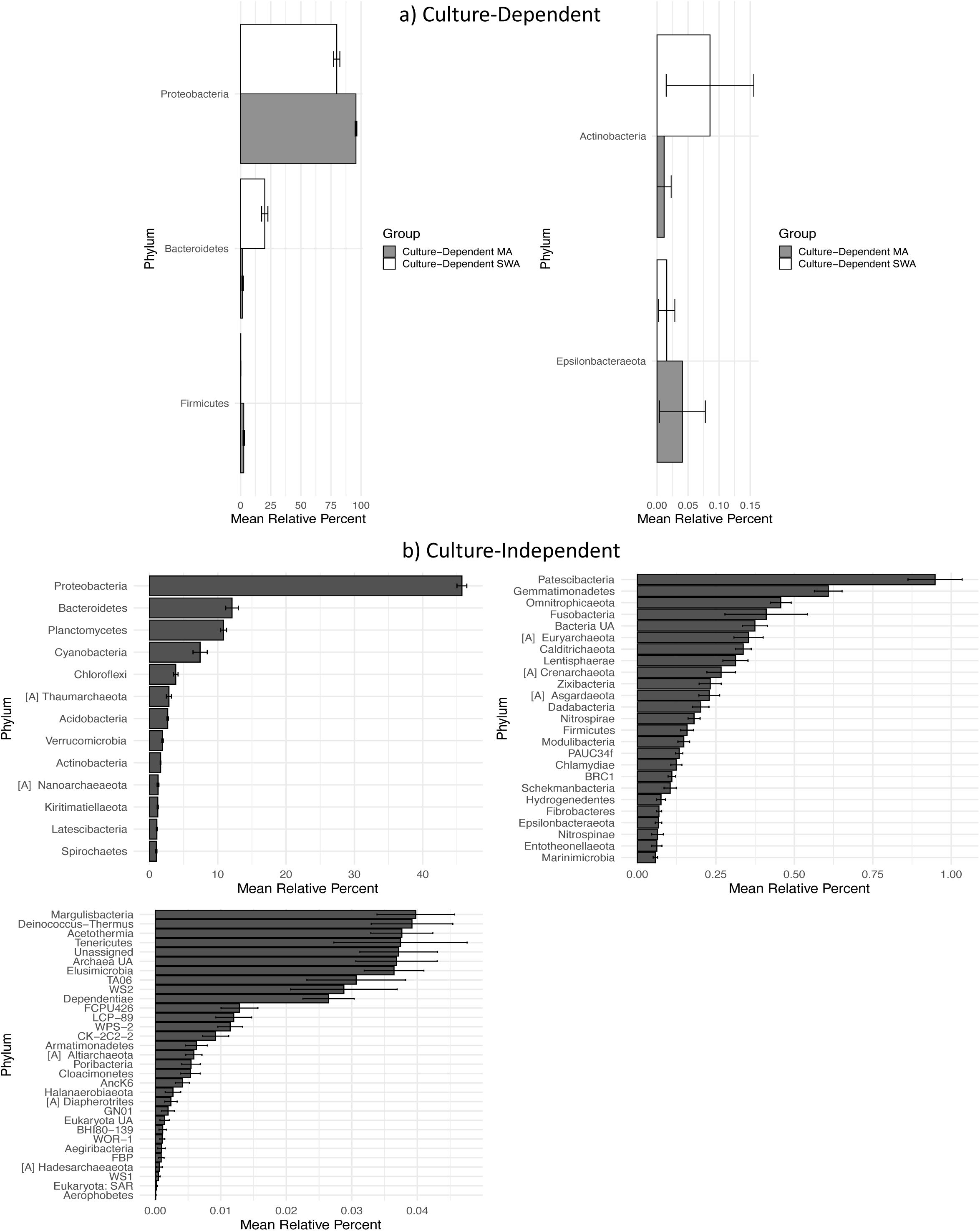
Rank abundance of phyla detected using a) culture-dependent methods and b) culture-independent methods. Light gray bars represent taxa associated with marine agar (MA) while white bare represent seawater agar (SWA) samples. Archaea are denoted as [A] and error bars indicate standard error.

At the genus level, the ASVs detected on SWA were assigned to 102 genera while those detected on MA were assigned to 76 genera (Table S1). CD methods identified 128 different genera, of which 50 were detected using both SWA and MA while 52 were unique to SWA and 26 were unique to MA (Figure 2). *Vibrio* represented the most abundant genus observed on both media, accounting for 27% and 44% of the sequences detected on SWA and MA, respectively. Other relatively abundant genera cultured included *Ruegeria* (SWA 25.4% & MA 35.1%), *Persicobacter* (SWA 18.8% & MA 0.7%), *Microbulbifer* (SWA 6.0% & MA 3.5%), and *Alteromonas* (SWA 3.9% & MA 1.7%) (Figure S3b). Most taxa detected with CD methods averaged <1% of the community, including 92 SWA assigned genera and 62 MA assigned genera (Table S1).

**Figure 2.**
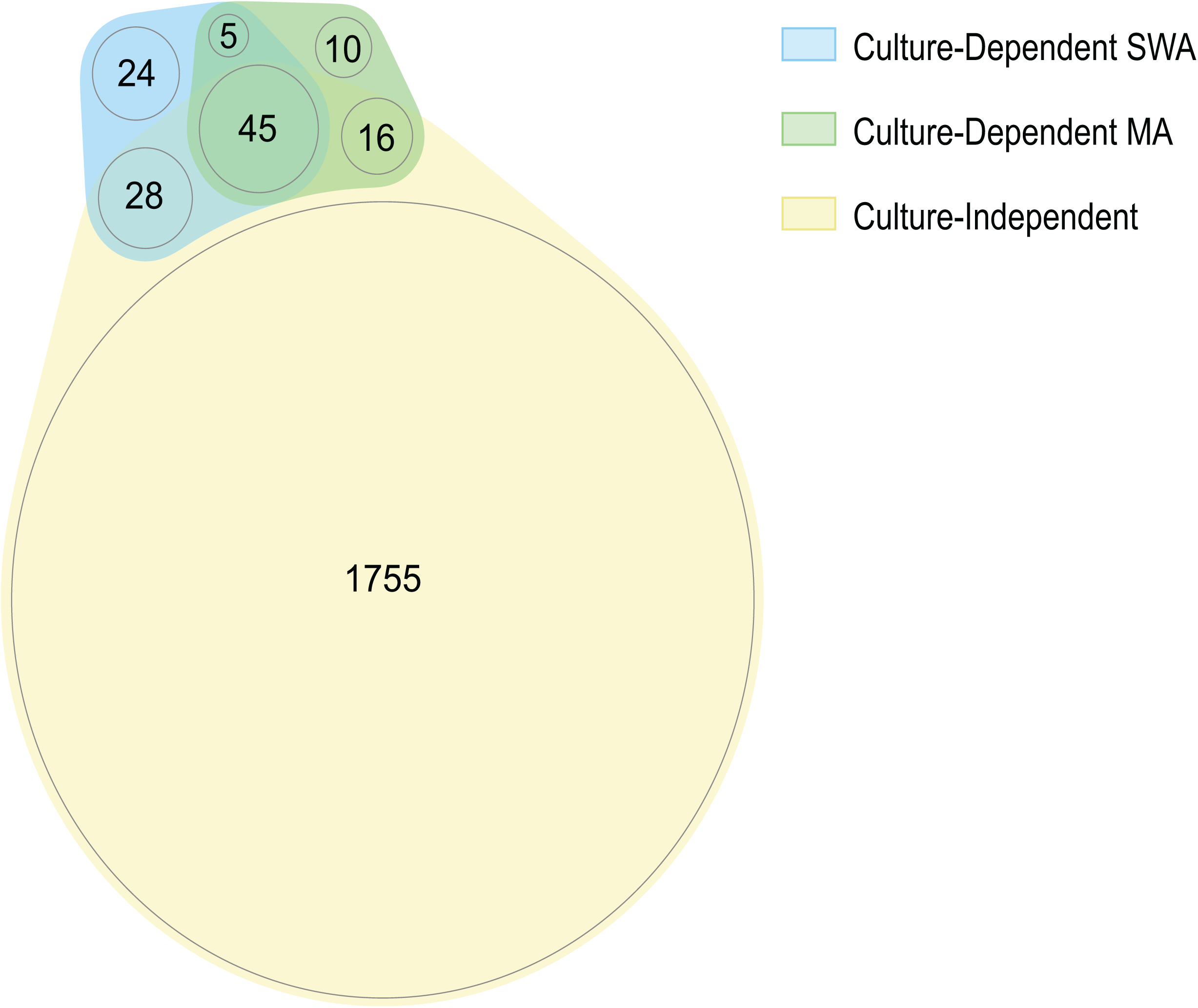
Proportionally scaled Venn diagram of microbial genera detected using culture-dependent (SWA and MA) and culture-independent methods.

### Culture-Independent (CI) Microbial Diversity

CI diversity for the same five sites was considerably greater than what was detected using the CD techniques, averaging 2,942 ± 133 ASVs (Figure S1b). CI communities included 68 phyla (57 Bacteria, 9 Archaea and 2 Eukarya) (Figure 1b, Figure S3a), with about half of the sequences assigned to Proteobacteria. After Proteobacteria, the relatively most abundant phyla were Bacteroidetes, Planctomycetes and Cyanobacteria. On average across all samples, 55 of the 68 phyla detected (81%) represented less than 1% of the relative community, and thus can be considered rare. When combined, these rare taxa averaged ∼7% of the total community, indicating their importance to community composition. More than 15 phyla represent either candidate phyla or unannotated taxa, indicating that microbial diversity at the phylum level remains poorly characterized in marine sediments (Figure 1b).

In total, 1,844 genera (1,728 Bacteria, 113 Archaea and 3 Eukarya) were observed using the CI technique (Figure 2), with the communities across all five sites displaying fairly rich (Figure S4a; Faith’s PD range 79.15-222.88) and even distributions (Figure S4b; Pielou’s evenness index range 0.81-0.93). The most commonly observed genus was *Woeseia*, which averaged 4.96% of the relative community across all five sites (Figure S3b). Of the 1,844 genera identified, only 16 averaged ≥1% of the community. Many CI ASVs were not annotated at the genus level or were annotated as “uncultured,” suggesting they belong to poorly described taxa. Those that averaged ≥1% of the community and could be identified at the genus level included *Woeseia* (4.96%), *Xenococcus* (2.49%), *Zeaxanthinibacter* (2.18%), Candidatus *Nitrosopumilus* (1.65%), *Pleurocapsa* (1.46%), *Chroococcidiopsis* (1.10%), and *Rhodopirellula* (1.07%) (Table S1). Minor (≤0.001 average relative percent), non-target amplification of eukaryotic sequences was observed. These were annotated as unassigned eukaryotic or ciliate associated (Table S1).

### Method Comparisons

In total, ASVs were annotated to 1,883 genera across all methods used in this study (Figure 2). As expected, the CI communities were significantly richer (Figure S2a; Faith’s PD Kruskal-Wallis H=52.076 p<0.001) and included 1,755 genera that were only detected using this approach (Figure 2). An additional 89 genera were detected using both the CI and CD approaches. Of those 89 genera, 45 were detected on both media types, 28 only on SWA and 16 only on MA (Figure 2). Surprisingly, 39 genera detected in culture were not detected using the CI technique (Figure 2; Table 1).

**Table 1.**
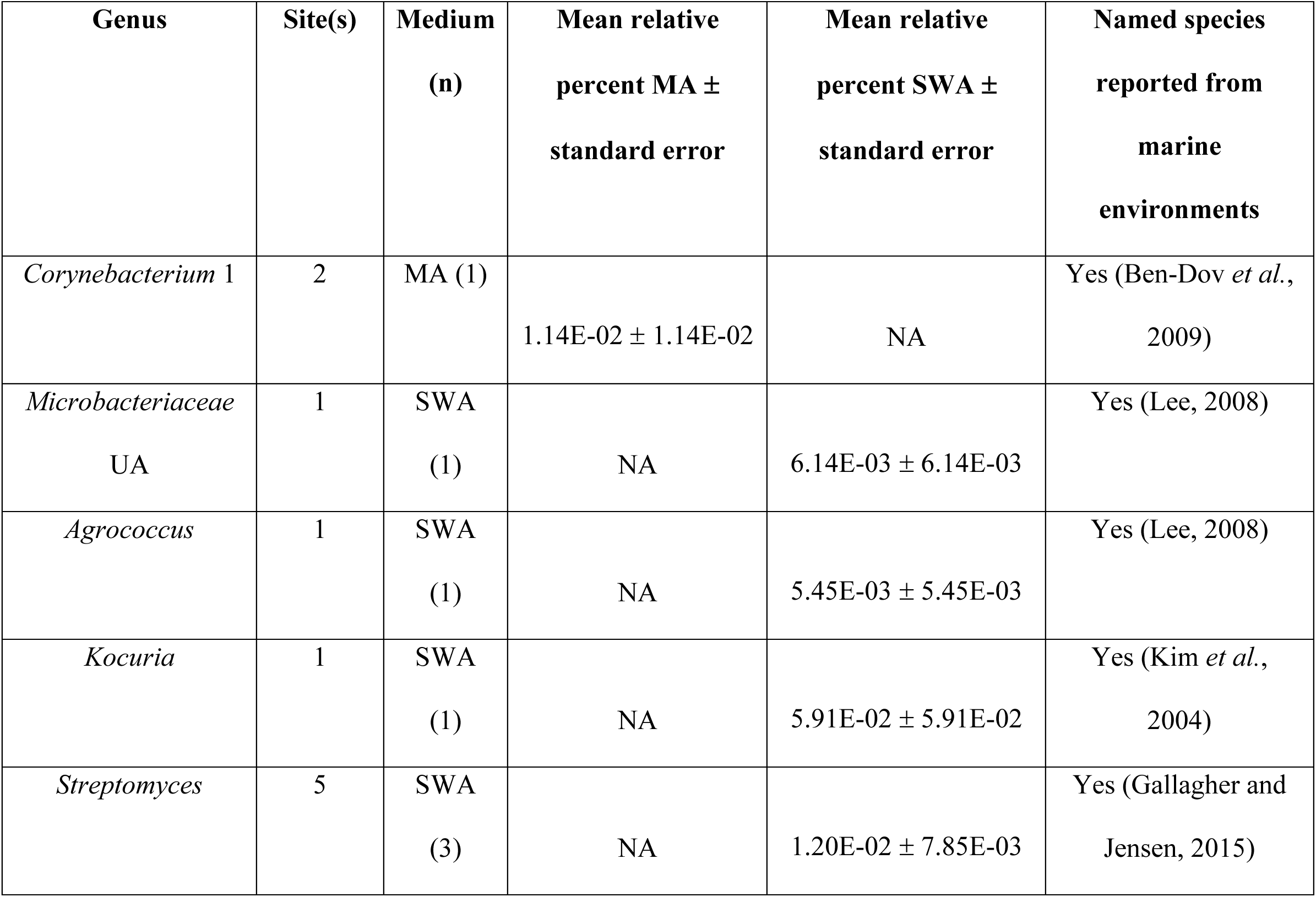
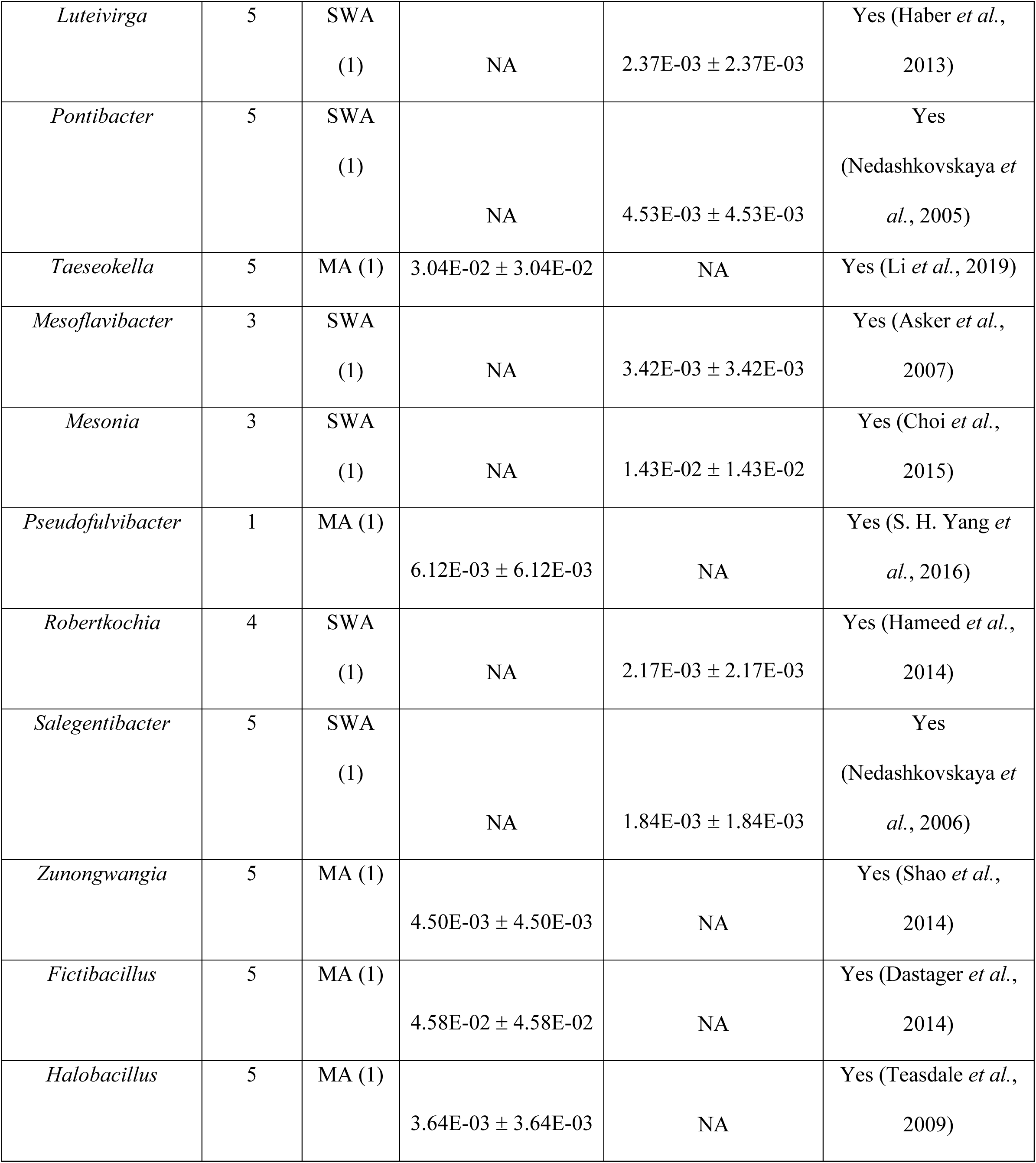
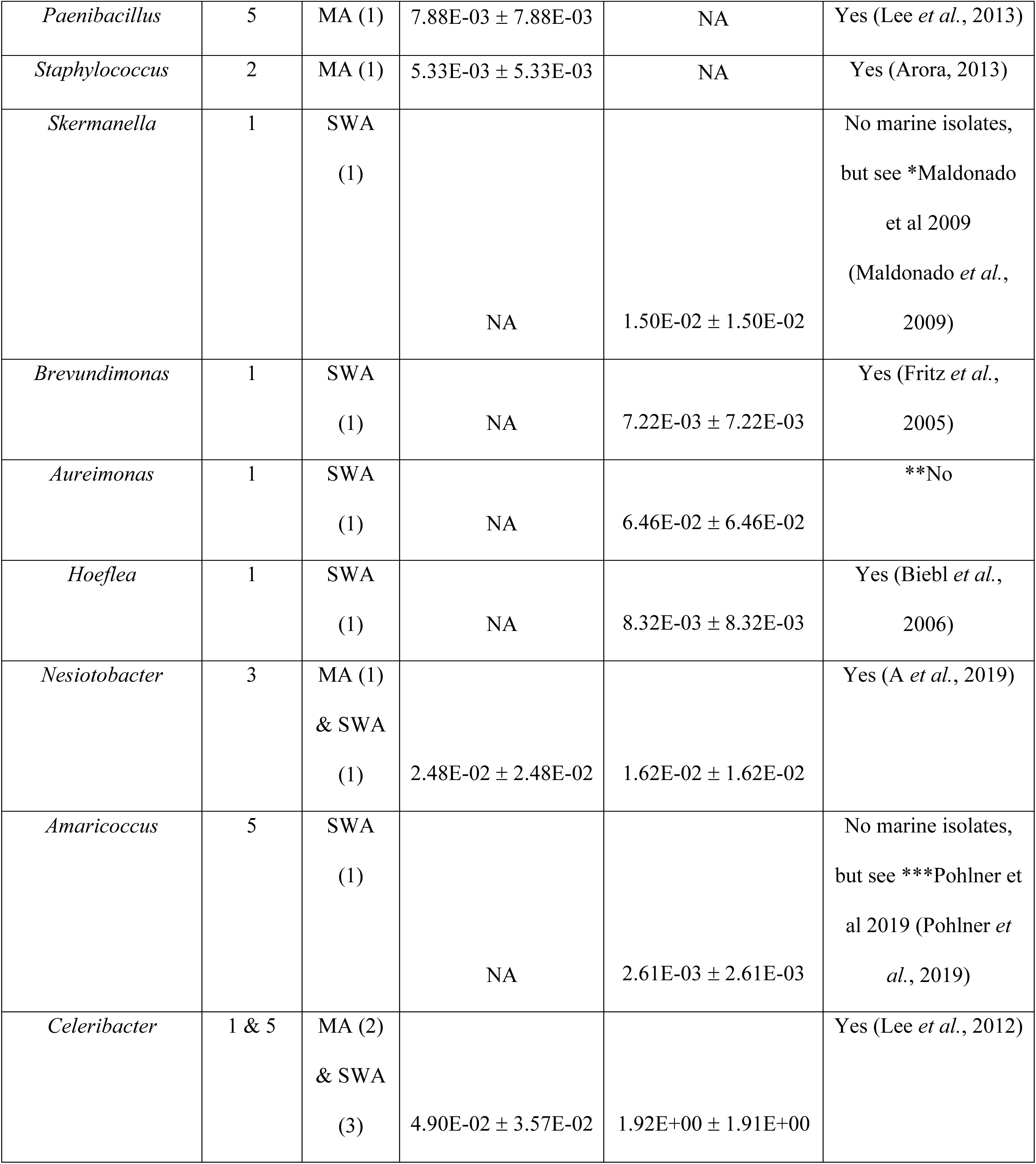
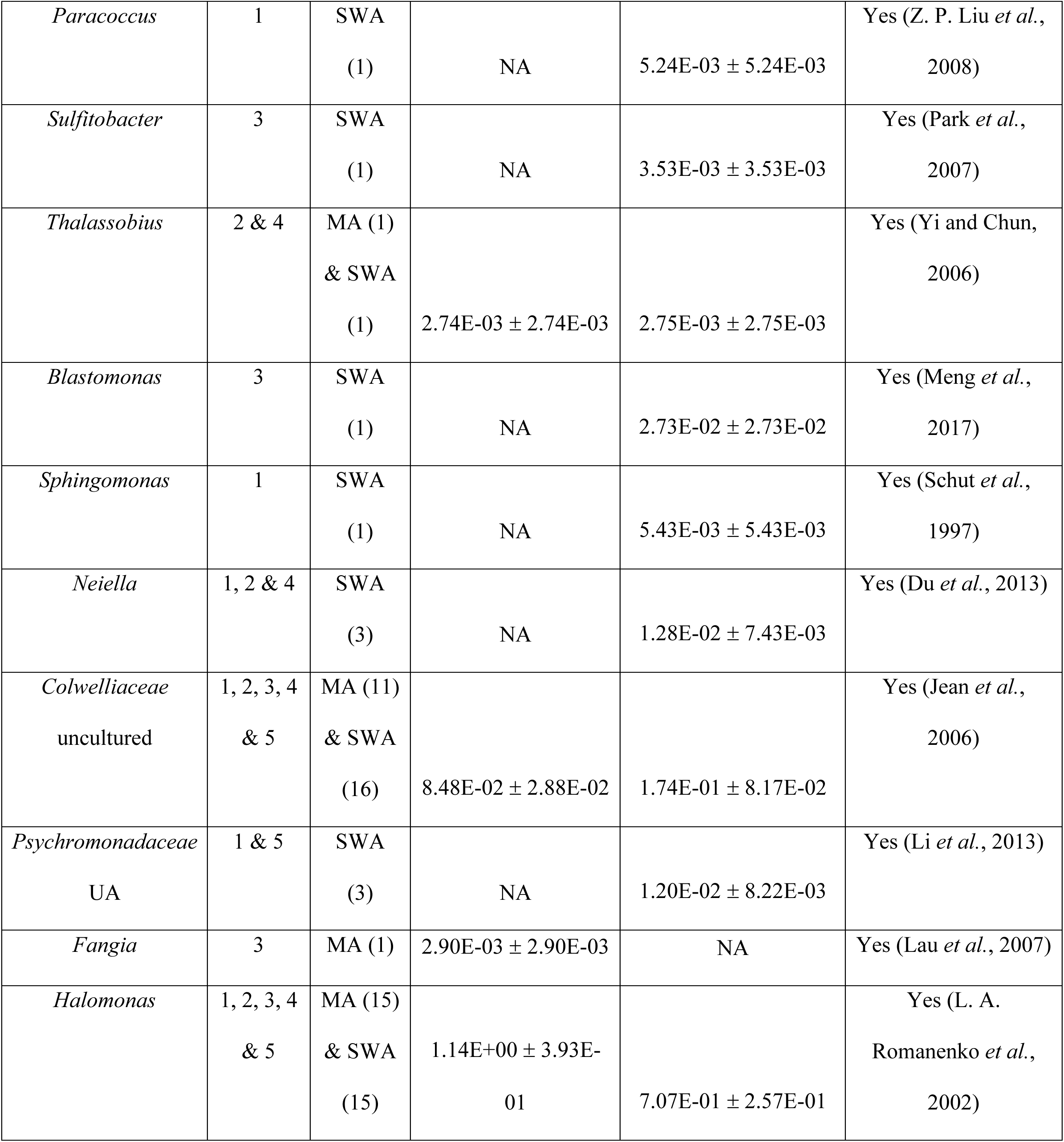
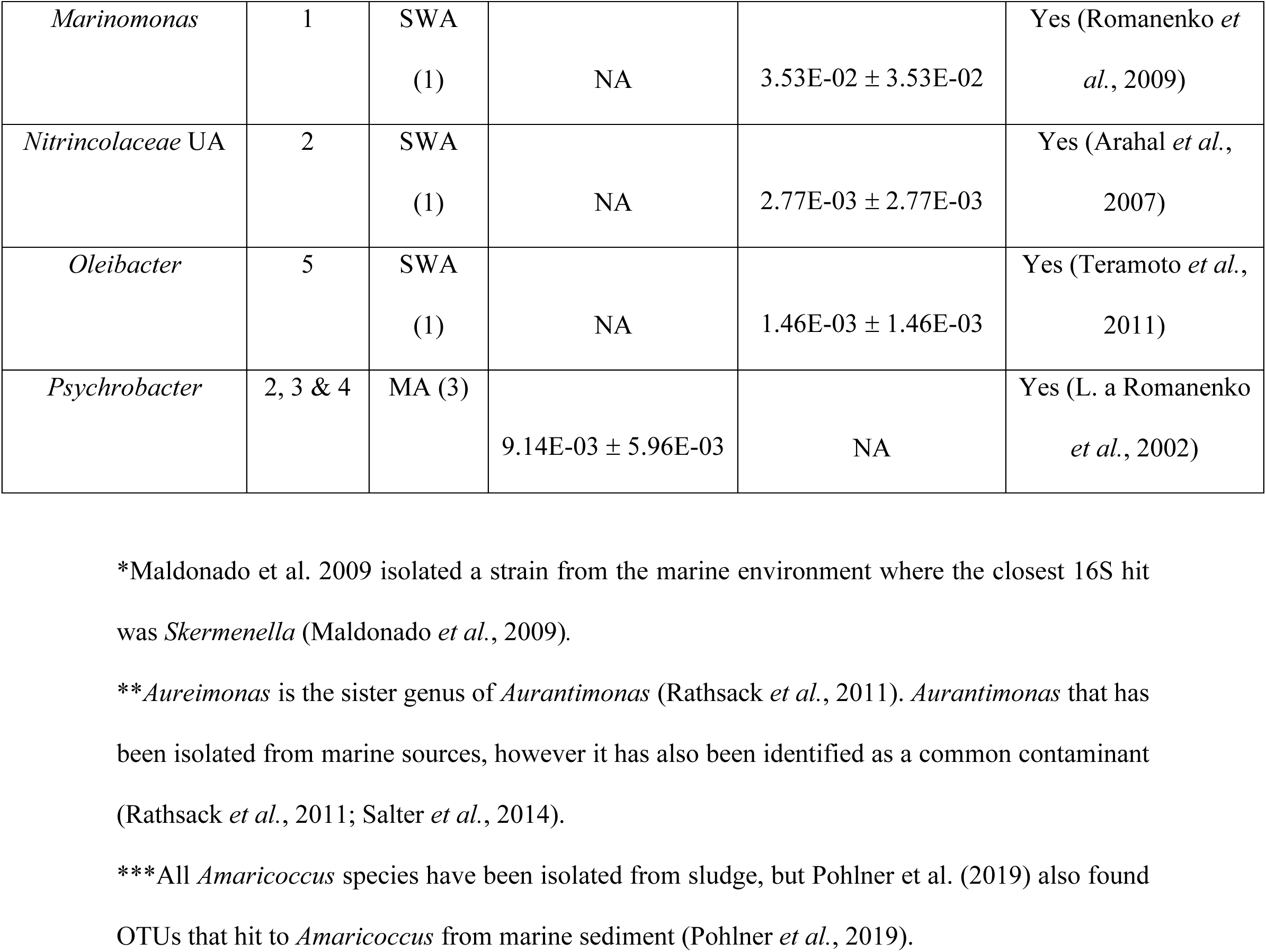
Amplicon sequence variants (ASVs) annotated to the genus level detected only with culture-dependent methods (n= number of detected replicates). Taxonomic assignments made with the SILVA v132 database through QIIME2-2020.2. If the genus was not annotated (UA), the lowest taxonomic rank is indicated. NA = not applicable. If the genus has named species from the marine environment, at least one example is cited.

Of the 39 genera that were only detected with CD methods, 24 of these genera were specific to SWA, ten were specific to MA, and five were detected on both SWA and MA (Figure 2). All of the genera uniquely detected using the CD technique were relatively rare, with 37 averaging <1% of the community (Table 1). The two most relatively abundant genera detected in culture but not using the CI technique were *Celeribacter*, which averaged 1.92% and 0.05% of the SWA and MA communities, respectively, and *Halomonas*, which averaged 0.71% and 1.14% of the SWA and MA communities, respectively. Taxa specific to the SWA method included Proteobacteria, Bacteroidetes and Actinobacteria such as *Marinomonas, Mesonia* and *Streptomyces*. Taxa only detected via MA cultures included members of the Firmicutes, Bacteroidetes and Proteobacteria such as *Fictibacillus, Taeseokella* and *Fangia* (Table 1). While the vast majority of taxa detected using CD methods have previously been isolated from marine sources, both the SWA and MA yielded genera that do not have named marine species. Interestingly, for all but one of these genera, closely related strains have been detected in marine samples (Table 1).

Not surprisingly, the CI approach identified numerous taxa with few or no cultured representatives. For example, the phylum Latescibacteria (aka WS3), which is commonly detected in CI studies (Youssef *et al*., 2015; Farag *et al*., 2017; Lloyd *et al*., 2018), and to the best of our knowledge does not have a cultured representative, was also identified as ca. 1.07% of the CI communities. Additionally, bacteria from the widely distributed and diverse phyla Acidobacteria, Patescibacteria and Gemmatimonadetes averaged 2.7%, 0.95, and 0.6% of the relative CI community respectively, but few strains from these phyla have been cultured (Hugenholtz *et al*., 2001; Ward *et al*., 2009; DeBruyn *et al*., 2011; Soro *et al*., 2014; Lemos *et al*., 2019).

Both CD and CI ASVs were also assessed in relation to their nearest cultured relative in the SILVA database (Figure 3, Figure S5). Not surprisingly, many of the ASVs shared greater similarity with sequences from other CI studies than with known type or cultured strains (Figure 3). ASVs from CD samples had a median similarity of 99.6% with cultured representatives, but there were a few distinct outliers (Figure 3, Figure S5). One ASV from CD samples was identified with SILVA as an uncultured Chitinophagales and shared only ∼79% similarity with a cultured representative. When that ASV was assessed with the NCBI BLAST database, the most similar sequence was a Muribaculaceae bacterium with ∼85% sequence similarity originally isolated from a pig gut. The other ASV from CD samples that was <90% similar to any culture in the SILVA database, matched with a recently cultured sponge microbe called *Xanthovirga aplysinae* (99.6% similar) (Goldberg *et al*., 2020). There were also a handful of sequences between 90-95% similar to cultured sequences. Subsequent searching with BLAST confirmed the lower similarity for some, while others had >99% similarity to recent isolates from marine sources such as holothurians and corals. In contrast to the CD samples, the median sequence similarity for CI samples was 85.4% (Figure 3, Figure S5). Many of the ASVs <60% similar were identified as archaea with SILVA. Of those, some shared greater similarity (∼80%) with members of the Microgenomates group of bacteria in the NCBI database.

**Figure 3.**
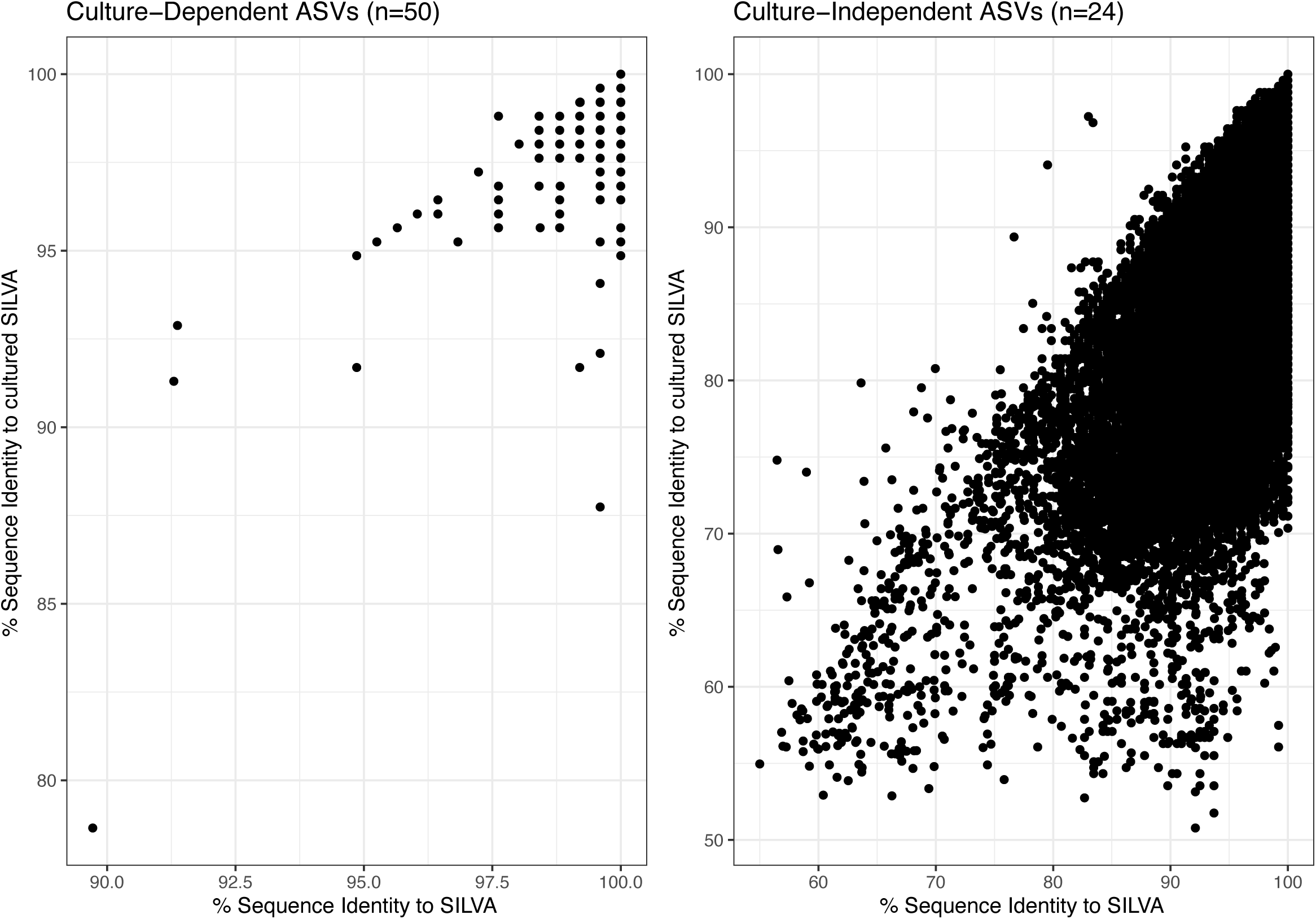
Scatterplot denoting the amplicon sequence variant (ASV) similarity from culture-dependent and culture-independent samples to the nearest cultured or type strain extracted from the SILVA v138.1 database (y-axis) versus the nearest sequence identified with the full non-redundant SILVA v138.1 database (x-axis).

When assessing beta diversity and the presence/absence of ASVs in each community, the CD communities were found to be significantly different than CI communities (Figure 4a; Unweighted UniFrac PERMANOVA pseudo-F=39.826, p=0.001). There was one CI replicate that was distinct from the rest of the CI samples (Figure 4), likely due to the large (>20%) fraction of an unknown genus of Flavobacteriales present within that sample. When considering sample site, cultured communities showed no pattern while CI communities seemed to cluster by site (Figure 4b). Weighted beta diversity analysis also showed significant separation between CD and CI communities (Figure 4c; Weighted UniFrac PERMANOVA pseudo-F=125.15, p=0.001) and between the nutrient rich (MA) and nutrient poor (SWA) media types (Figure 4c). Since there appeared to be clustering based on sites within the CI communities (Figure 4b & d), only CI samples were considered for subsequent CI community visualizations and comparisons.

**Figure 4.**
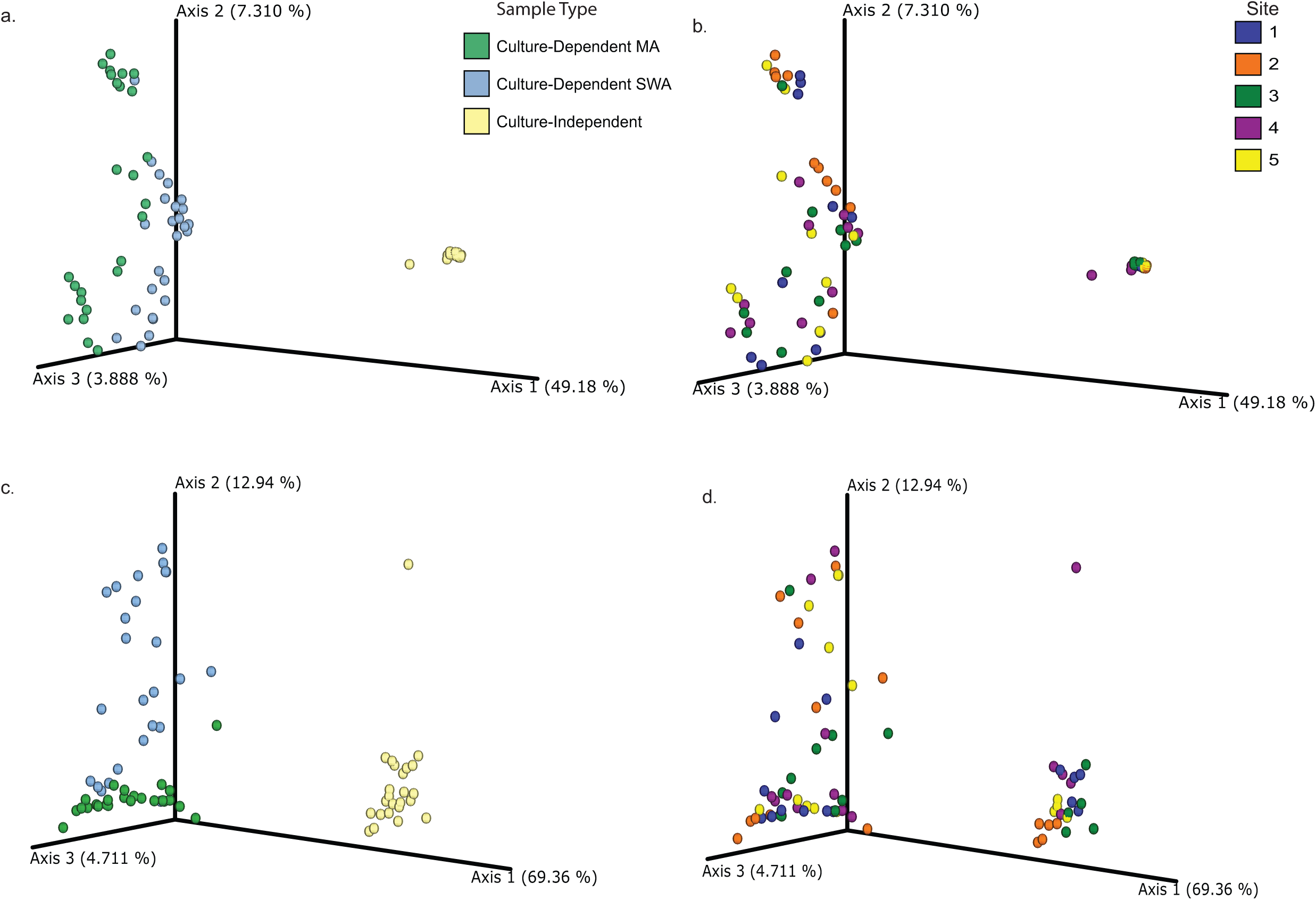
UniFrac analysis of culture-independent and culture-dependent sediment microbial communities visualized via principle coordinates analysis (PCoA). Unweighted results colored by a) sample type and b) site. Weighted results colored by c) sample type and d) site.

### Culture-Independent (CI) Site Comparisons

The five sites sampled were within 5 km of each other (Figure S6, Table S2), but represented different habitats: an 8 m deep spur and groove reef (site 1), a 20 m deep reef slope (site 2), a 1 m deep sand patch in a seagrass bed at the mouth of a mangrove island (site 3), a 6 m deep seamount in a lagoon (site 4), and a 1 m deep sand and rubble patch at the marine station’s dock (site 5). Both weighted and unweighted UniFrac analyses on the CI communities indicate a significant difference among site communities (Figure 5; Unweighted UniFrac PERMANOVA pseudo-F=2.982, p=0.001; Weighted UniFrac PERMANOVA pseudo-F=7.987, p=0.001).

**Figure 5.**
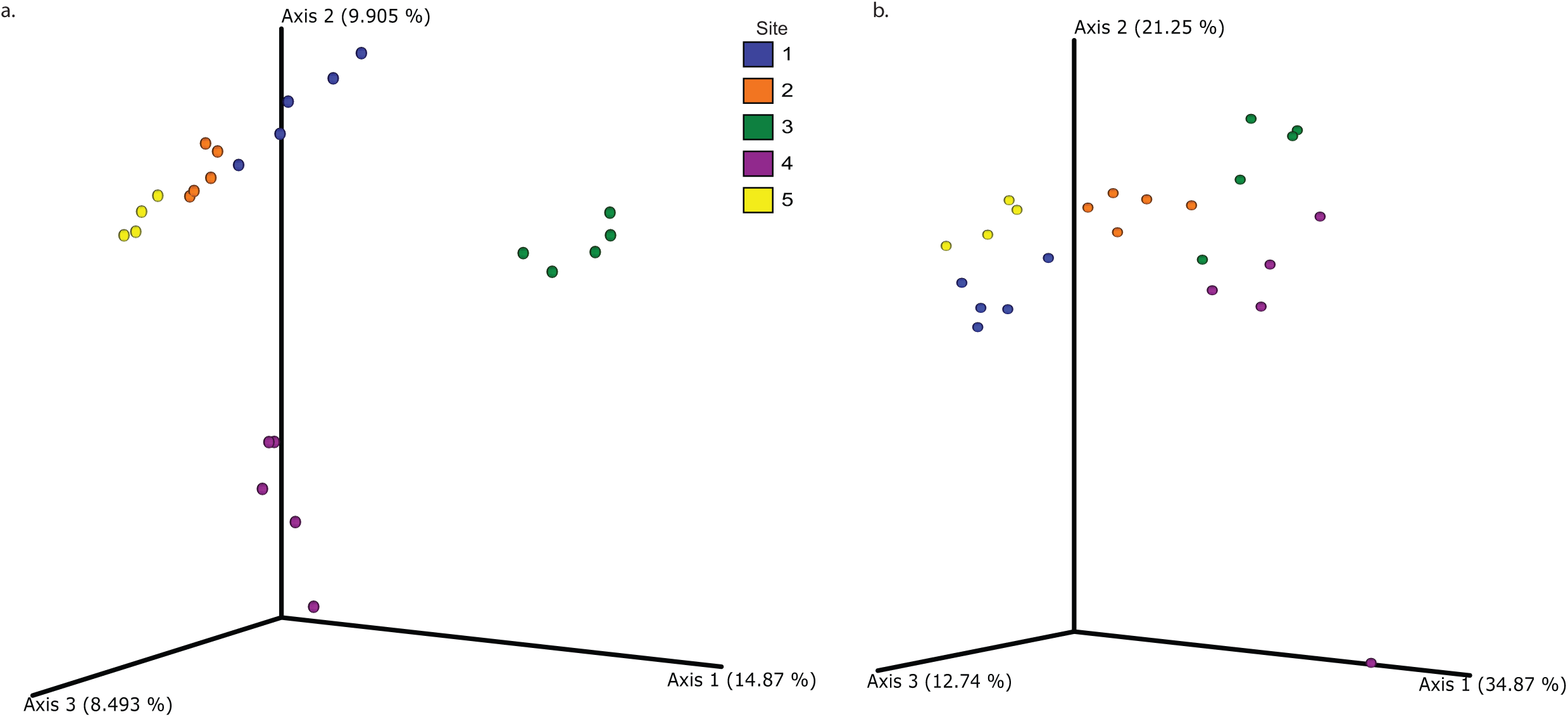
Unifrac analysis of culture-independent sediment microbial communities across five sites. a) Unweighted UniFrac and b) weighted UniFrac. Communities visualized via principle coordinates analysis (PCoA). PERMANOVA results indicate sites are significantly different from each other in both unweighted (pseudo-F=2.982, p=0.001) and weighted (pseudo-F=7.987, p=0.001) analyses.

Site 3 appeared the most distinct from the other communities in the unweighted UniFrac (Figure 5a) and showed higher richness than the four other sites (Figure S4a). Some of the taxonomic differences between sites included relatively more Proteobacteria at site 2, relatively fewer Cyanobacteria at sites 3 and 4, and about twice the relative amount of Chloroflexi at site 3 when compared to the other sites (Figure S3a). Additionally, an analysis of composition of microbiomes (ANCOM) done at the genus level identified seven taxa that were significantly different by site, six of which were related to site 3. These were identified as *Marixanthomonas*, an unidentified genus of the BD2-7 (Family Cellvibrionales), an uncultured gammaproteobacterium that was relatively more abundant at site 3, and two taxa that were absent from site 3 (*Stanieria*, an unknown genus of Xenococcaceae, and un uncultured MBAE14 gammaproteobacterium).

### Culturing Efficiency

We assessed culturing efficiency in two ways. First, we quantified percent efficiency by determining the total number of ASVs cultured vs. the number identified using CI techniques. The alpha diversity rarefaction curves based on observed ASVs revealed that saturation was achieved using both techniques (Figure S1), thus ensuring that effective comparisons could be made. The number of cultured ASVs represented 1.82% and 1.33% (for SWA and MA, respectively) of the ASVs detected using the CI approach. Second, we calculated the overlap in taxonomic assignment from domain down to ASV using the two approaches (Table 2, Figure S7). Based on taxonomic assignment, 3.95% of the genera detected using the CI approach were also detected in culture on SWA medium while 3.31% were detected on MA. Thus, while the amount of diversity cultured will vary with the method used, our results generally exceeded the <1% value typically reported in association with the great plate count anomaly (Staley and Konopka, 1985; Martiny, 2019; Steen *et al*., 2019).

**Table 2.** Percent cultured across taxonomic levels. Percent cultured calculations were based on taxonomic overlap in culture-dependent (CD) and culture-independent (CI) samples. Percent taxa unique to CD samples represents the number of taxa only identified with CD methods as a proportion of the total taxa identified at the corresponding level.

|  | **Percent Cultured** |  | **Percent Taxa Unique to Culture-Dependent Samples** |  |
| --- | --- | --- | --- | --- |
| **Taxonomic Level** | **SWA** | **MA** | **SWA** | **MA** |
| Domain | 33.33 | 33.33 | 0 | 0 |
| Phylum | 7.35 | 7.35 | 0 | 0 |
| Class | 2.39 | 3.19 | 0 | 0 |
| Order | 2.91 | 2.75 | 0.46 | 0.15 |
| Family | 3.36 | 2.76 | 0.51 | 0.34 |
| Genus | 3.95 | 3.31 | 1.54 | 0.80 |
| Species | 3.16 | 2.64 | 1.77 | 1.22 |
| ASV | 0.39 | 0.33 | 0.86 | 0.68 |

## Discussion

It is widely recognized that CD approaches are inadequate for measuring microbial diversity. Early comparisons of colony counts to cell counts indicated that <1% of environmental bacteria were cultured, a phenomenon referred to as the “great plate count anomaly” (Razumov, 1932; Staley and Konopka, 1985). The great plate count anomaly was the first indication that a majority of environmental bacteria were not readily cultured using standard techniques, which has subsequently driven numerous studies seeking to improve culturability (Kaeberlein *et al*., 2002; Tamaki *et al*., 2009; Tanaka *et al*., 2014; Rygaard *et al*., 2017). The advent of sequence-based CI methods brought the extent of uncultured microbial diversity into better perspective (Lynch and Neufeld, 2015; Hug *et al*., 2016; Lloyd *et al*., 2018). But while high-throughput amplicon sequencing methods, such as 16S rRNA gene surveys, are valuable tools for assessing microbial communities, there are biases and limitations with these methods. For instance, DNA extraction (Brooks *et al*., 2015), primer selection (Fischer *et al*., 2016; Laursen *et al*., 2017; Wear *et al*., 2018; Willis *et al*., 2019), PCR amplification (Brooks *et al*., 2015) and template concentration (Kennedy *et al*., 2014) have all been shown to impact microbial community profiles and the use of the 16S rRNA gene as a taxonomic marker can miss up to 10% of environmental sequences (Eloe-Fadrosh *et al*., 2016). Additionally, variables used for sequence analysis, such as operational taxonomic unit clustering vs. denoising affect diversity estimates (Patin *et al*., 2013; Callahan *et al*., 2017; Edgar, 2017, 2018; Nearing *et al*., 2018; Straub *et al*., 2019) and resulting data interpretations are further complicated by the compositional nature of amplicon data (Gloor *et al*., 2017).

Given advances in both CD and CI techniques, it is surprising though that few studies have sought to reassess bacterial diversity estimates using both approaches (but see some example comparisons in river sediment: (Pédron *et al*., 2020), lake sediment: (Elfeki *et al*., 2018), seawater: (Rygaard *et al*., 2017), cheese: (Perin *et al*., 2017), and lungs: (Dickson *et al*., 2014)). Here, we used CD and CI techniques to explore the microbial communities in marine sediments and estimate culturing efficiency by sequencing bacteria directly from agar plates as opposed to the more traditional approach of isolating or counting colonies. The 16S rRNA gene sequences amplified from sediment eDNA required ∼50,000 reads, post quality control and denoising, to approach saturation in the alpha rarefaction curves. This indicates that relatively deep sequencing is needed to capture the microbial diversity present in these complex communities. A recent study comparing global diversity of marine sediments detected over 34,000 ASVs associated with bacteria and archaea across 299 sites (Hoshino *et al*., 2020). In total, our study found over 27,000 ASVs from CD and CI samples around Carrie Bow Cay, Belize. One major difference between the global study (Hoshino *et al*., 2020) and ours is sequencing depth. While Hoshino et al. (2020) were able to do an impressive global analysis on a large number of samples, none of their rarefaction curves reached saturation, suggesting that much of the rare biosphere was missed. Given the richness levels detected at fine scales in sediment (Probandt *et al*., 2018) and that the majority of taxa identified in our sediments were in low relative abundance (<1%), deep sequencing is needed to detect ecologically relevant members of the rare biosphere and to fully profile sediment communities (Lynch and Neufeld, 2015; Jousset *et al*., 2017).

As expected, CD richness estimates were considerably lower, averaging 54 and 39 ASVs per sample for seawater agar (SWA) and marine agar (MA) respectively, compared to >2,000 ASVs per CI sample. While we did not perform colony counts, the number of ASVs detected appeared higher than the number of colonies readily visualized by eye on the plates after eight days of incubation, suggesting that much of the diversity detected may have arisen from micro-colonies that could not be easily counted or isolated using standard practices. This result likely helps account for the observation that culturing efficiency, as defined by the number of detected ASVs, on SWA and MA relative to the CI results was 1.82 and 1.33%, respectively, or 3.15% in total. Given that only two cultivation media were used, these results support a re-evaluation of the 1% culturability paradigm. It would also be beneficial to perform colony counts in parallel to sequencing plates to help delineate methodological differences in assessing the great plate count anomaly and to confirm the presence of micro-colonies in culture. Pairwise approaches have been successfully used in some studies (e.g. Perin *et al*., 2017; Rygaard *et al*., 2017; Elfeki *et al*., 2018; Pédron *et al*., 2020), but not yet in marine sediments to our knowledge. Culturability varies depending on the ecosystem and complexity of the community, so additional studies across systems and with varying methodologies will be beneficial for proper re-assessment of the great plate count anomaly.

We were also interested in assessing culturing efficiency based on the taxa detected in culture compared to CI samples. While, we determine the taxonomic overlap between CD and CI methods from domain to ASV, we focused on the genus level given limitations with species assignment when using short regions of the 16S rRNA gene (Liu *et al*., 2008; Větrovský and Baldrian, 2013; Yang *et al*., 2016; Johnson *et al*., 2019). When assessing culturing efficiency based on the genera detected with both CD and CI methods, culturability increased to 3.95 and 3.31% on SWA and MA respectively (and increased to 5.42 and 4.04% if including genera only identified with CD methods). While this highlights that cultivation may in fact be more effective than the original experiments of the great plate count anomaly, the lack of marine sediment representatives in culture was apparent given the high frequency at which our taxonomy was denoted as “uncultured” or unassigned. While we would expect culturability to increase with taxonomic level, surprisingly we found that the culturing percentage remained fairly consistent (∼3%) until the phylum (∼7%) and domain level (∼33% since no Archaeal or Eukaryotic sequences were detected in culture). This finding supports previous research documenting the limited knowledge on marine sediment microbial diversity (Baker *et al*., 2021).

While we further explored differences in communities at the genus level, it is important to note that there is considerable intra-genus diversity in microbes (Cordero *et al*., 2012; Patin *et al*., 2015), thus limiting the conclusions we can draw from studies such as ours based on a short fragment of the 16S rRNA gene. Additionally, given functional variation within microbial lineages, assessing communities at different taxonomic levels would provide differing insight. For instance, Hoshino *et al*. (2020) evaluated ASV differences in relation to the phylum level which can provide some insight into broad functional patterns (e.g. aerobic vs anaerobic metabolism) but lacks the resolution and/or associated metagenomic data to concretely answer functional questions. Additionally, taxonomic assignment might vary based on the reference database. For instance, SILVA may only identify an ASV to the order level while a closer NCBI match would provide taxonomy to the genus level. Moreover, if taxonomy is annotated to the same level, reference databased can disagree (Pollock *et al*., 2018). Thus, it can be difficult to extract ecological meaning from taxonomic community profiles, especially in understudied environments like sediments that lack cultured representatives and genomic data for many lineages (Baker *et al*., 2021).

Here, we defined cultured taxa as those that were identified after applying a background threshold to remove any ASVs that had fewer than 70 reads in their corresponding samples. The three cultured phyla that occurred in >1% relative abundance were Proteobacteria, Bacteroidetes and Firmicutes. Among these, Proteobacteria also represented the largest fraction of the CI community, indicating that members of the most abundant phylum detected can be readily cultured. While both Bacteroidetes and Firmicutes were detected on SWA and MA, SWA had a considerably larger fraction of Bacteroidetes (∼20% compared to ∼2%) while MA had more Firmicutes than SWA (∼3% compared to ∼0.1%), demonstrating how these different media enrich for different taxa. Additionally, the cultivation of significantly greater microbial richness on the relatively nutrient poor medium SWA supports previous results (Watve *et al*., 2020) and suggests that, even for sediment communities, high nutrient concentrations can be inhibitory. Interestingly, the candidate phylum PAUC34f was detected in every replicate of our CI analyses, yet this phylum remains uncultured to date (Chen *et al*., 2020). While PAUC34f was not detected in our post-filtering CD data, it was detected in one SWA replicate prior to correcting for the inoculum control. Given that dilution to extinction in a nutrient poor medium led to the cultivation of the ubiquitous SAR11 clade of marine bacterioplankton (Rappé *et al*., 2002), our decision to pool eDNA obtained from the dilutions prior to sequencing may have masked evidence for the cultivation of this phylum.

Approximately 20% of the ASVs detected using the CD method could only be assigned to broad groups, such as the Roseobacter clade CHAB-I-5, or remained unclassified at the genus level. Given the accuracy of removing erroneous sequences while obtaining greater taxonomic resolution with denoising pipelines (Callahan *et al*., 2016, 2017; Amir *et al*., 2017; Nearing *et al*., 2018; Prodan *et al*., 2020), our ASVs without taxonomic assignment likely represent undescribed taxa obtained in culture. Our subtraction of inoculum controls further supports this suggestion. In addition to culturing taxa that could not be assigned to a known genus, we also cultured genera that contained few named species. For instance, the genus *Ascidiaceihabitans* only contains one species, isolated from the tunicate *Halocynthia aurantium* (Kim *et al*., 2014), yet was detected on six SWA replicates (max ∼27% relative percent) and three MA replicates (max ∼2% relative percent). We also observed the genus *Endozoicomonas* in two replicates of MA (max ∼2% relative percent). *Endozoicomonas* bacteria have been found in association with a wide variety of marine organisms including corals (Bayer *et al*., 2013), tunicates (Schreiber *et al*., 2016) and sea slugs (Kurahashi and Yokota, 2007). A recent CI study revealed the potential for diverse functional roles for *Endozoicomonas* as a symbiont while also postulating that this bacterium has a free-living stage based on its large genome size (Neave *et al*., 2017). Our detection of *Endozoicomonas* in multiple replicates potentially represents a previously uncultured, free-living stage of this taxon.

In addition to separation of the CD vs CI communities observed in beta diversity analyses, we also saw clustering by site in the CI samples, with site 3, a sand patch near a seagrass bed and a mangrove island, seemingly the most distinct. Gribben et al. 2017 found that seagrass sediment communities can reduce the success of an invasive macrophyte, indicating their importance for the overall health of these ecosystems (Gribben *et al*., 2017). ANCOM results identified seven taxa at the genus level that significantly differed between sites, with three taxa found to have the highest relative abundance at site 3 and three taxa completely absent from site 3. Further studies are needed to determine what, if any, role these taxa may play in seagrass communities and what physical and chemical characteristics may play a role in structuring these communities. Changes in clustering patterns of CI communities were also observed when using unweighted vs weighted UniFrac. The unweighted UniFrac showed clearer separation across sites in the CI communities. Given that weighted UniFrac accounts for the relative abundances of taxa, this finding suggests that there are commonalities in the dominant community members across the five sites. Additionally, all significant genera identified through ANCOM were rare members of the community, with the maximum relative percent abundance of ∼0.2%. Therefore, it appears that rare members of the community are likely driving site differentiation in beta diversity. While we cannot determine functional differences in our sites with this study, previous research has demonstrated the disproportionate role rare microbes may play in communities (Jousset *et al*., 2017; Bech *et al*., 2020). Given that the vast majority of taxa identified in our study represented <1% of the relative communities at each site, it would be valuable to explore how rare microbes may functionally impact sediment communities in future work.

One surprising result was the detection of cultured taxa that were not detected using CI methods. Similar results were reported from Mediterranean water samples (Crespo *et al*., 2016) and recent work in a freshwater system (Pédron *et al*., 2020). Almost all genera only detected with CD methods would likely be considered members of the rare biosphere. Eight of the genera detected only in CD samples have been previously identified as laboratory contaminants including *Corynebacterium, Kocuria, Paenibacillus, Brevundimonas, Hoeflea, Paracoccus, Sphingomonas, Acinetobacter,* and *Psychrobacter* (Salter *et al*., 2014), however they all also have named species from the marine environment (Table 1). Given the identification of marine representatives and our filtering methods, we believe these taxa to be members of our communities rather than contaminants. To the best of our knowledge, *Aureimonas* is the only genus detected exclusively from CD methods that has not been previously reported from marine samples. It was detected on one MA replicate from Site 1 and represented 1.62% of that community. *Aureimonas* is a sister genus to *Aurantimonas*, which does include marine representatives (Rathsack *et al*., 2011), suggesting that the ASV might have been misannotated due to high sequence similarity. Alternatively, this may be the first report of the genus *Aureimonas* from the marine environment. Importantly, the CD methods revealed ASVs that could not be identified to the genus level, or were simply annotated as “uncultured”, suggesting that new taxa had been cultured using these relatively simple approaches (Table 2, Table S1).

One of the potential limitations in using environmental enrichment methods is the difficulty in isolating pure cultures. Bacteria may fail traditional isolation attempts for a variety of reasons including obligate associations with co-occurring microbes or metabolic needs that are not met when the strain is isolated from the community. Thus the use of new techniques such as reverse genomics (Cross *et al*., 2019) and metagenomics in conjunction with traditional methods can aid in culturing efforts (Lewis *et al*., 2020). Additionally, innovative culturing techniques that utilize environmental conditions such as diffusion chambers (Bollmann *et al*., 2007) and the ichip (Berdy *et al*., 2017) also provide a means to increase culturing success. It would be interesting to further interrogate the unique taxa we detected in our culturing approach to determine if they could be isolated using traditional methods such as dilution to extinction, traditional methods with altered medium (e.g. Rygaard *et al*., 2017) or if new techniques might prove more fruitful. One of the divergent strains detected in CD samples was subsequently identified as *Xanthovirga aplysinae.* Given the recent description of this genus, isolated with traditional methods on MA (Goldberg *et al*., 2020), it is likely that continued culturing efforts in underexplored environments will yield novel diversity. It is also possible that our threshold for defining culturability was not stringent enough due to differences between sample and control sediments, in which case further efforts should focus on taxa that were detected in relatively higher abundances across multiple cultures.

In conclusion, the sediment microbial communities analyzed here were highly diverse, with the majority of genera representing rare (<1%) members of the community. The unique detection of taxa using CD methods supports the value of these techniques in conjunction with CI methods to assess community composition. Additionally, it may be valuable to further explore variation in microbial communities across spatial scales to ensure adequate sampling and diversity assessments.

## Experimental Procedures

### Sample Collection and Processing

In September 2015, divers collected marine sediment samples from five locations around the Smithsonian Field Station at Carrie Bow Cay, Belize (Figure S6; Table S2). At each site, five replicate Whirl-Pak® (Nasco) bags were filled with sediment from a 3 m^2^ area. Upon return to the field station, 20 ml of wet sediment from each Whirl-Pak® was transferred into 50 ml falcon tubes with 20 ml of RNAlater® and stored at 4 °C before transport on dry ice to Scripps Institution of Oceanography (SIO) where they were stored at −40 °C prior to DNA extraction. Two types of media prepared at SIO were used for on-site, culture-dependent sample processing: seawater agar (SWA) comprised of 16 g agar and 1 liter natural seawater, and 50% marine agar (MA) comprised of 0.5 g yeast extract, 2.5 g peptone, 16 g agar, and 1 liter natural seawater. Both media contained the antifungal agent cyclohexamide at a final concentration of 200 μg/ml. Freshly collected sediment samples were diluted 1:2 with autoclaved seawater in 4 ml vials, vigorously shaken, and further diluted 1:10 and 1:100 after which 50 μl of each dilution (1:2 1:10, 1:100) was inoculated onto each medium, spread with a sterile glass rod, and the plates allowed to dry in a laminar flow hood then sealed with parafilm. This resulted in a total of 150 plates (3 dilutions x 5 replicate sediments x 2 media x 5 locations), which were transported back to SIO. In an effort to facilitate colony growth but prevent one or two bacteria from swamping the plates, we kept the plates at room temperature for eight days post inoculation and then stored them at −40 °C prior to DNA extraction.

Two sets of control samples for the CD method were generated in January 2020. To control for DNA contamination present in the media or reagents, plates made with both of the experimental medium types were inoculated with a known amount (∼3×10^7^ cells) of *Vibrio coralliilyticus,* then immediately parafilmed and stored at −40 °C. To assess the approximate read counts that might be expected from the initial inoculation without subsequent colony growth, dilution series of local sediment (San Diego, CA) were plated following the experimental plate methods. Three replicate dilution series were plated for each medium, each was spiked with a known amount (∼3×10^7^ cells) of *Vibrio coralliilyticus* to ensure adequate DNA concentrations for sequencing, parafilmed immediately to prevent colony growth, and stored at −40 °C.

### DNA Extraction

Environmental DNA (eDNA) was extracted from approximately 1 g of freshly thawed sediment per sample following physical (bead beating) and chemical (phenol-chloroform) DNA extraction methods (Patin *et al*., 2013). One replicate from site 5 was lost resulting in a total of 24 sediment samples processed for CI analysis. DNA extractions were performed in duplicate for each sediment sample (2 g of sediment in total extracted per sample) and the extracts combined prior to purification. For the CD analyses, agar plates were left to thaw at room temperature for 30 min and 3 ml molecular grade water was added to the surface. A heat sterilized metal loop was used to scrape the surface of each plate and the resulting suspension pipetted into a 15 ml falcon tube. The three dilutions plated for each sediment were combined into a single falcon tube and centrifuged at 8,000 RPM (9,803 RCF) and 4 °C for 5 minutes generating 50 samples (5 replicate sediments x 2 media x 5 locations). The supernatant was removed and the bottom 2 ml including the cell pellet were distributed into two ceramic bead-beating tubes prior to DNA extraction following the protocol applied to the sediments (Patin *et al*., 2013). All control samples were extracted using the same protocol described above, however due to samples being lost in transit to the sequencing facility, minimal DNA remained for subsequent sequencing. As a result, replicates for each control type were pooled, thus producing four control samples: a SWA blank control spiked with *V*. *coralliilyticus,* a SWA inoculum control spiked with *V. coralliilyticus,* a MA blank control spiked with *V*. *coralliilyticus,* and a MA inoculum control spiked with *V. coralliilyticus*.

### PCR and Sequencing

The v4 region of the 16S rRNA gene was PCR amplified using the primers 515F (TCGTCGGCAGCGTCAGATGTGTATAAGAGACAG GTGYCAGCMGCCGCGGTAA) and 806Rb (GTCTCGTGGGCTCGGAGATGTGTATAAGAGACAGGGACTACNVGGGTWTC TAAT) (Caporaso *et al*., 2012). PCR was performed following the Phusion Hot Start Flex 2x Master Mix protocol with an annealing temperature of 60 °C and 1 μl of 5 ng/μl DNA. Products were cleaned using ExoSap-IT® before adding Nextera XT (Illumina) indices and sequencing adapters with the following PCR program: 98 °C for 1 min followed by five cycles of 98 °C for 10 sec, 65 °C for 20 sec and 72 °C for 30 sec with a final extension at 72 °C for 2 min. Gel electrophoresis was used to confirm the presence of a PCR product of the predicted size. Sequences were normalized based on DNA concentration, pooled and cleaned with AMPure XP beads. The purified library was then sent to the Institute for Genomic Medicine (IGM), University of California, San Diego (UCSD) for sequencing on an Illumina MiSeq v2 500 cycle at a depth of 130k reads per sample. For controls, DNA was sent to Novogene (South Plainfield, NJ) for library preparation following their proprietary methods and sequencing with the above described primers (515F & 806Rb) on an Illumina NovasSeq.

### Analysis

Raw sequences were imported into QIIME2-2020.2 (Bolyen *et al*., 2018) and denoised using the DADA2 (Callahan *et al*., 2016) denoise-paired pipeline with an input of p-trim-left-f of 19 and p-trim-left-r of 20 to remove primers. Based on the raw files, p-trunc-len-f and p-trunc-len-r were set to 250 and 155 base pairs respectively and chimeras were removed with the default consensus method. In an effort to control for background inoculum that remained present but was not actively growing, all amplicon sequence variants (ASVs) associated with the controls were quantified to their relative read abundances (Table S3). After accounting for ASVs associated with *Vibrio*, the most abundant ASV present was identified as an unknown bacterium with 67 reads in one of the control inoculum samples (Table S3). Since all other ASVs associated with the inoculum controls, outside of *Vibrio* which was used as a spike-in, contained less than 67 reads, we opted to set a minimum read threshold of 70 reads per feature for each CD sample before proceeding with subsequent analyses. This threshold removed approximately 50% of ASVs associated with CD samples (Table S4). Additionally, we checked each ASV associated with the four control samples and determined that none of the ASVs remained in the experimental samples after applying the filtration step. Taxonomy was then assigned using the SILVA v132 database (Quast *et al*., 2013) and samples were subsequently filtered to remove chloroplast and mitochondria sequences (Table S4) in QIIME2-2020.2.

In order to assess how similar ASV sequences were to cultured representatives, an approach based on Steen *et al*. (2019) was employed. Using the align.seqs command in Mothur (Schloss and Westcott, 2011), both CD ASVs and CI ASVs were aligned with sequences from the SILVA database. To compare to cultured strains, type strains [T] and cultured s[C] strains were searched for in SILVA and downloaded as an aligned fasta file including gaps. Since SILVA v138.1 was the database used for sequence extraction, the corresponding non-redundant full library was also downloaded (SILVA_138.1_SSURef_NR99_tax_silva_full_align_trunc.fasta). The align.report files were then filtered to remove any sequences with inadequate alignments (pairwise alignment lengths <250). Histograms based on similarity to the nearest cultured representative and scatterplots with the nearest cultured relative compared to the most similar sequence in the full library were generated using ggplots2 (Wickham, 2016) in R. Some of the ASVs were then further interrogated by using the NCBI BLAST tool by selecting to exclude uncultured sequences.

QIIME2-2020.2 was used to perform alpha and beta diversity analyses at the ASV level after rarefying samples to a depth of 62,830 reads. Associated statistical analyses were also performed with QIIME2-2020.2 at the ASV level using the non-parametric Kruskal-Wallis test (Kruskal and Wallis, 1952) for method and site comparisons in relation to alpha diversity indices (Faith’s Phylogenetic Diversity and Pielou’s Evenness) and multivariate PERMANOVA tests with 999 permutations (Anderson, 2001) for beta diversity (both weighted and unweighted UniFrac (Lozupone and Knight, 2005)) comparisons across methods and sites. ANCOM analysis (Mandal *et al*., 2015), which was specifically designed to address compositional microbial data, was performed at the genus level using QIIME2-2020.2 to determine genera that significantly differed across sites in culture-independent samples. Figures were generated using QIIME2-2020.2, RStudio version 3.6.2 (R Core Team, 2019), ggplot2 (Wickham, 2016), nVennR (Quesada, 2020), and Excel version 16.36.

### Data Availability

All raw sequences files are available through NCBI’s Sequence Read Archive (SRA). Accession numbers for CI files are SAMN08824420 – SAMN0882443, CD files are SAMN15932210 – SAMN15932259 and controls are SAMN15932260 – SAMN15932263.

## Acknowledgments

We would like to thank Robert Tuttle for assisting in the collection of samples and Gregory Amos and Alexander Chase for bioinformatic advice. Additionally, we would like to acknowledge Jessica Blanton and Eric Allen for providing sequencing indices and for assistance in library construction. We would also like to acknowledge the Smithsonian’s Carrie Bow Cay Field Station. This publication is a contribution to the Caribbean Coral Reef Ecosystems (CCRE) Program. Sequencing was conducted at the IGM Genomics Center, University of California, San Diego, La Jolla, CA.

This research was supported by the National Science Foundation Grant No. OCE-1235142 and based upon work supported by the National Science Foundation Graduate Research Fellowship under Grant No. DGE-1650112 and the Scripps Fellowship to A.M.D.

Any opinion, findings, and conclusions or recommendations expressed in this material are those of the authors(s) and do not necessarily reflect the views of the National Science Foundation. The authors declare no conflicts of interest.

## Supplemental Tables & Figures

**Table S1.** All genera assignments from QIIME2-2020.2 analysis with the SILVA v132 Database and the average relative percent of the community identified to each genus across CD and CI methods.

**Table S2.** Site information for samples collected around Carrie Bow Cay, Belize.

**Table S3.** Amplicon sequence variants (ASVs) detected in control samples sorted by decreasing frequency. Blank controls were marine agar (MA) or seawater agar (SWA) plates spiked only with Vibrio. Inoculum controls were MA or SWA plates spiked with Vibrio and fresh sediment inoculum. All controls were done in triplicate, but DNA was pooled prior to 16S sequencing resulting in one replicate per control sample type. ASVs were assigned after denoising with DADA2. Taxonomy was determined with SILVA v132. ASVs associated with the Vibrio spike-in were omitted from the table. See methods section for further details.

**Table S4.** Number of amplicon sequence variants (ASVs) remaining after filtration steps were applied. Sample types include the control blanks which were culture-dependent (CD) plates spiked with Vibrio, control with inoculum were CD plates spiked with Vibrio and fresh sediment inoculum, CD marine agar (MA) cultures, CD seawater agar (SWA) cultures and culture-independent samples. ASVs were generated from denoising with DADA2 and taxonomy was assigned with SILVA v132. All steps were performed in QIIME2-2020.2.

**Figure S1.** Mean alpha rarefaction curves across sediment samples from five sites in Belize. a) Culture-dependent results obtained using seawater agar (SWA) and marine agar (MA) media and b) Culture-independent results. Error bars represent standard error among replicates.

**Figure S2.** Alpha diversity boxplots of marine sediment microbial communities from Carrie Bow Cay, Belize in culture-dependent and culture-independent samples determined using a) Faith’s Phylogenetic Diversity Index and b) Pielou’s Evenness. Culture-dependent methods include the use of marine agar medium (MA) and seawater agar medium (SWA. Data points are overlayed on the boxplot to show variation.

**Figure S3.** Relative abundance of microbial communities in marine sediments. a) Phylum level culture-dependent diversity on two growth media (MA and SWA, left) and culture-independent diversity (right). b) Genus level culture-dependent diversity on two growth media (MA and SWA, left) and culture-independent diversity (right). Legends lists a) all phyla and b) the top 50 genera in order from most to least abundant and six rare genera that had a notable percentage in at least one replicate. Note that bar colors repeat for some rare taxa.

**Figure S4.** Culture-independent alpha diversity boxplot of marine sediment microbial communities from across five sites around Carrie Bow Cay, Belize determined using a) Faith’s Phylogenetic Diversity Index and b) Pielou’s Evenness.

**Figure S5.** Histograms of amplicon sequence variant (ASV) from culture-dependent and culture-independent samples and their similarity to previously cultured strains. Cultured representatives included both type and cultured strains extracted from SILVA v138.1.

**Figure S6.** Sediment sample collection site information.

**Figure S7.** Proportional Venn Diagrams denoting the number of taxa detected with culture-dependent and culture-independent methods (marine agar and seawater agar) across taxonomic levels.

