## Supplemental Figures for "Microbial diversity in tropical marine sediments assessed using culture-dependent and culture-independent techniques"

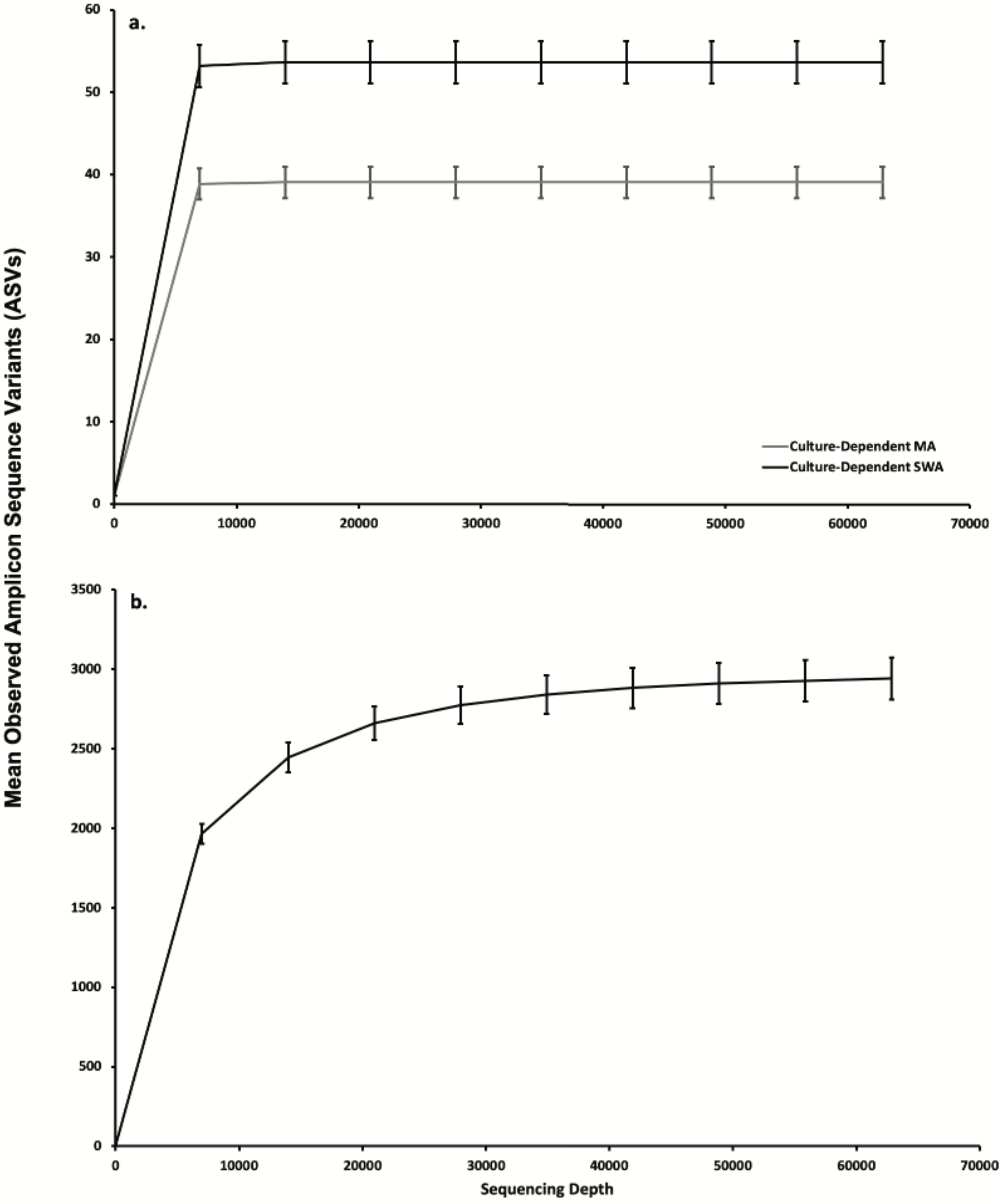


Figure S1. Mean alpha rarefaction curves across sediment samples from five sites in Belize. a) Culture-depend results obtained using seawater agar (SWA) and marine agar (MA) media and b) Culture- independent results. bars represent standard error among replicates.

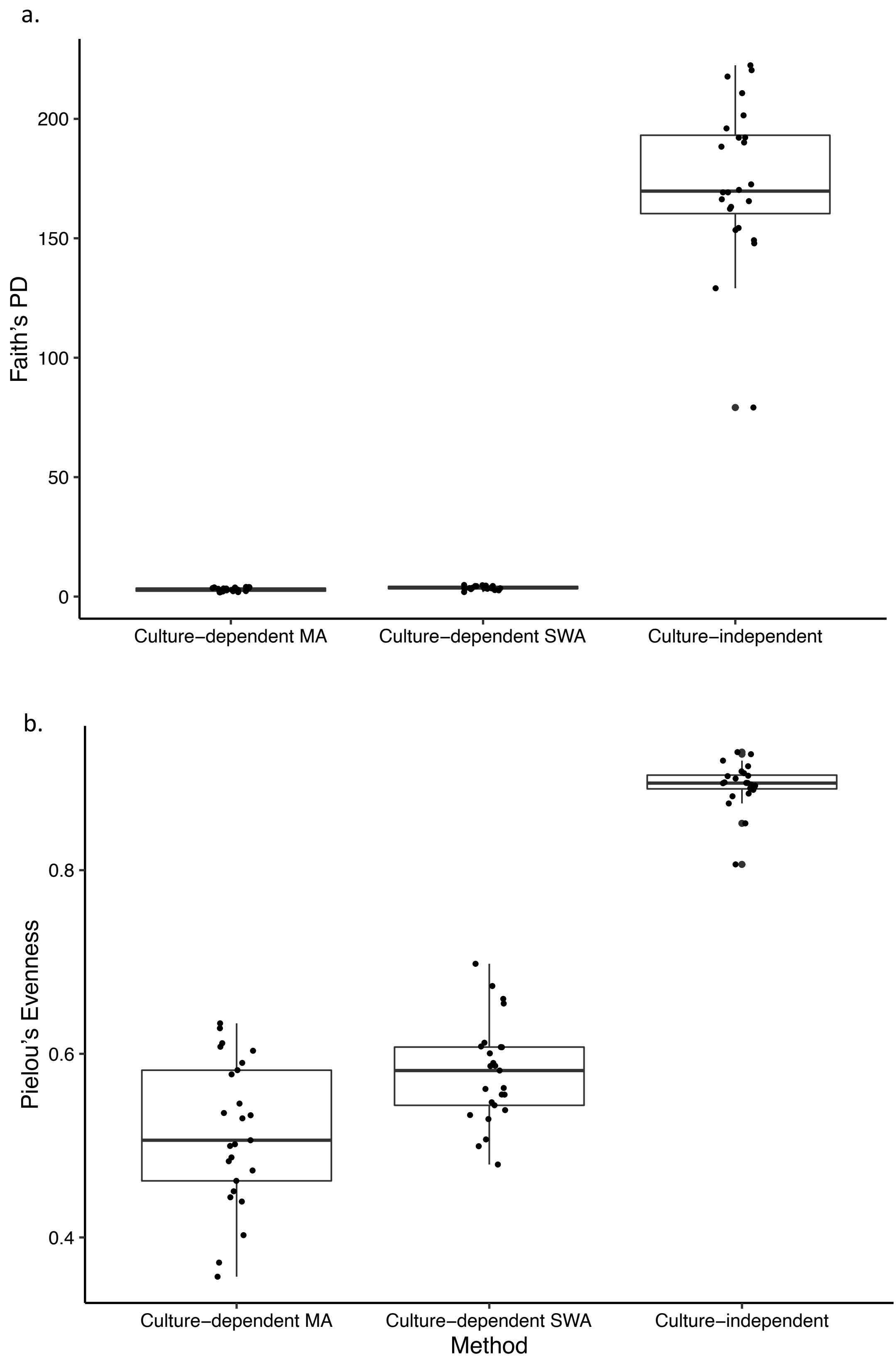


Figure S2. Alpha diversity boxplots of marine sediment microbial communities from Carrie Bow Cay, Belize in culture-dependent and culture-independent samples determined using a) Faith's Phylogenetic Diversity Index and b) Pielou's Evenness. Culture-dependent methods include the use of marine agar medium (MA) and seawater agar medium (SWA. Data points are overlayed on the boxplot to show variation.

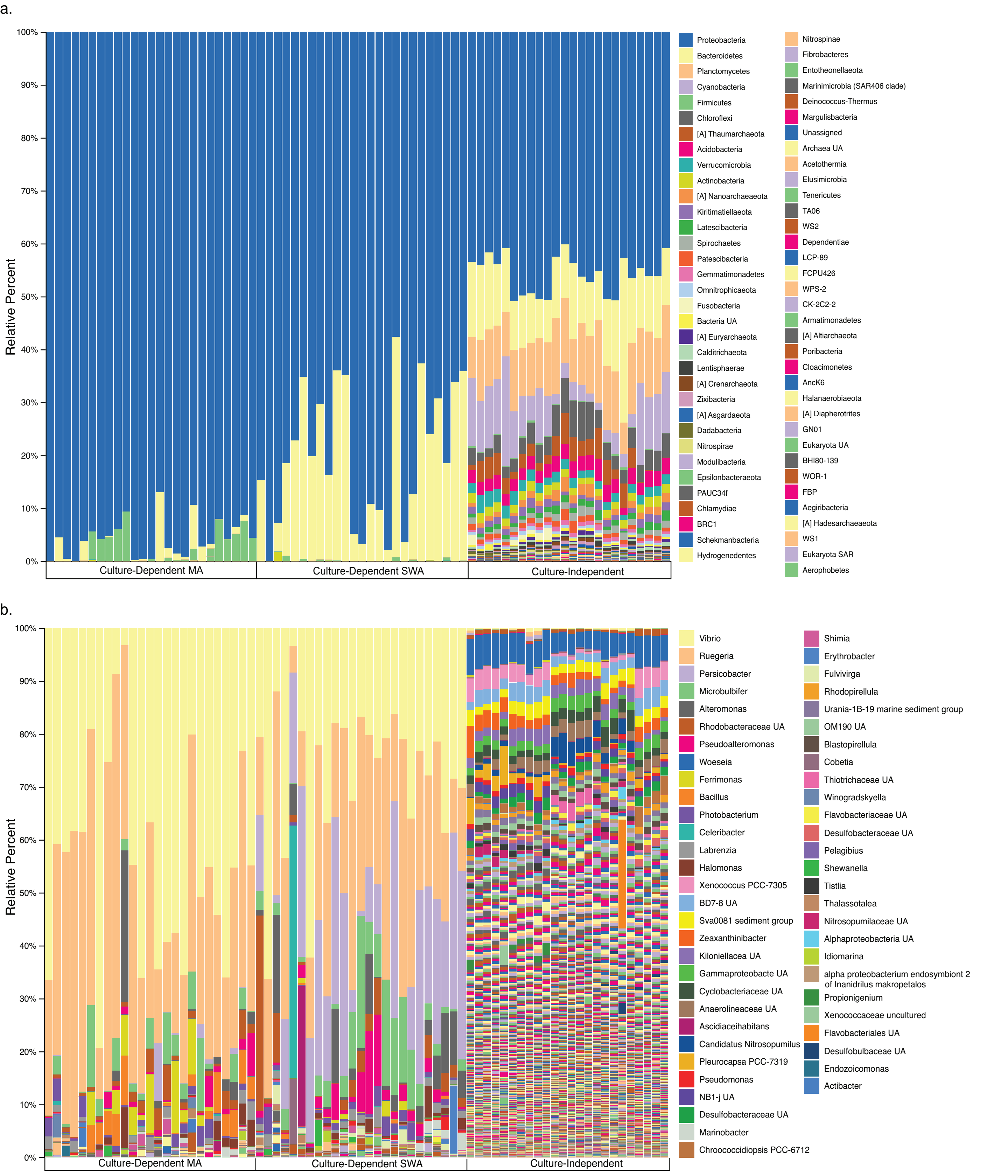


Figure S3. Relative abundance of microbial communities in marine sediments. a) Phylum level culture-dependent diversity on two growth media (MA and SWA, left) and culture-independent diversity (right). b) Genus level culture-dependent diversity on two growth media (MA and SWA, left) and culture-independent diversity (right). Legends lists a) all phyla and b) the top 50 genera in order from most to least abundant and six rare genera that had a notable percentage in at least one replicate. Note that bar colors repeat for some rare taxa.

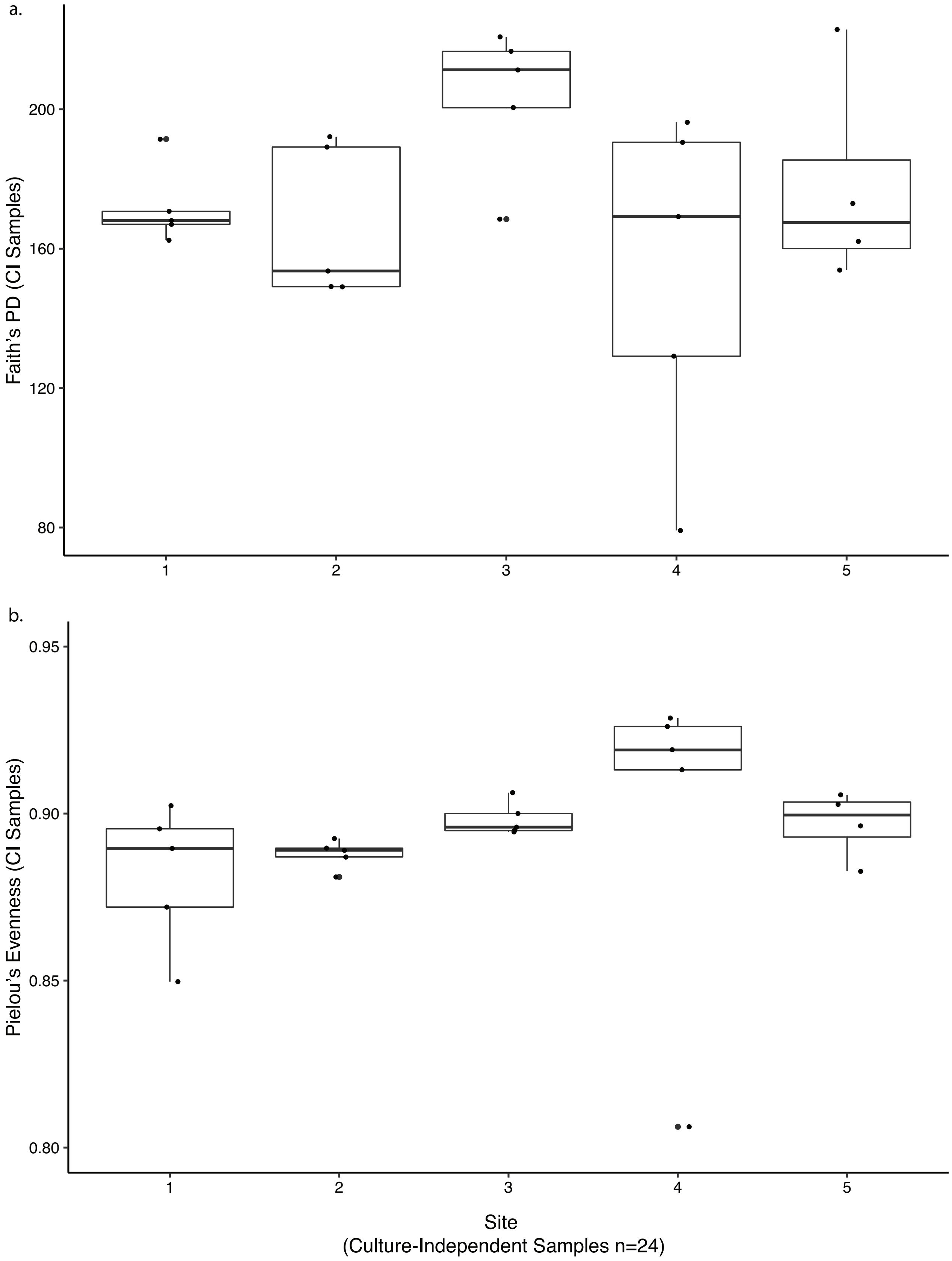


Figure S4. Culture-independent alpha diversity boxplots of marine sediment microbial communities from across five sites around Carrie Bow Cay, Belize determined using a) Faith's Phylogenetic Diversity Index and b) Pielou's Evenness.

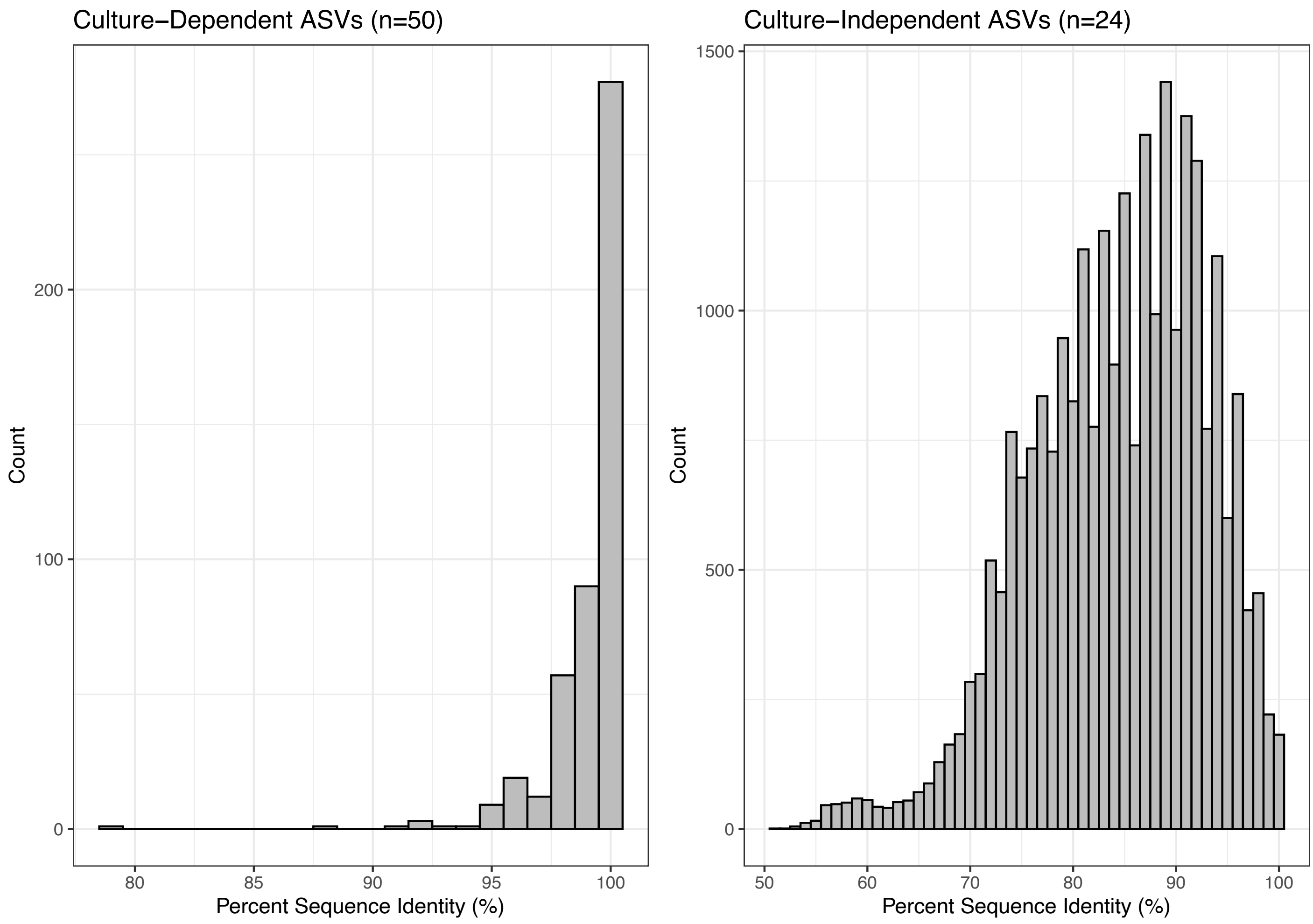


Figure S5. Histograms of amplicon sequence variant (ASV) from culture-dependent and culture-independent samples and their similarity to previously cultured strains. Cultured representatives included both type and cultured strains extracted from SILVA v138.1.

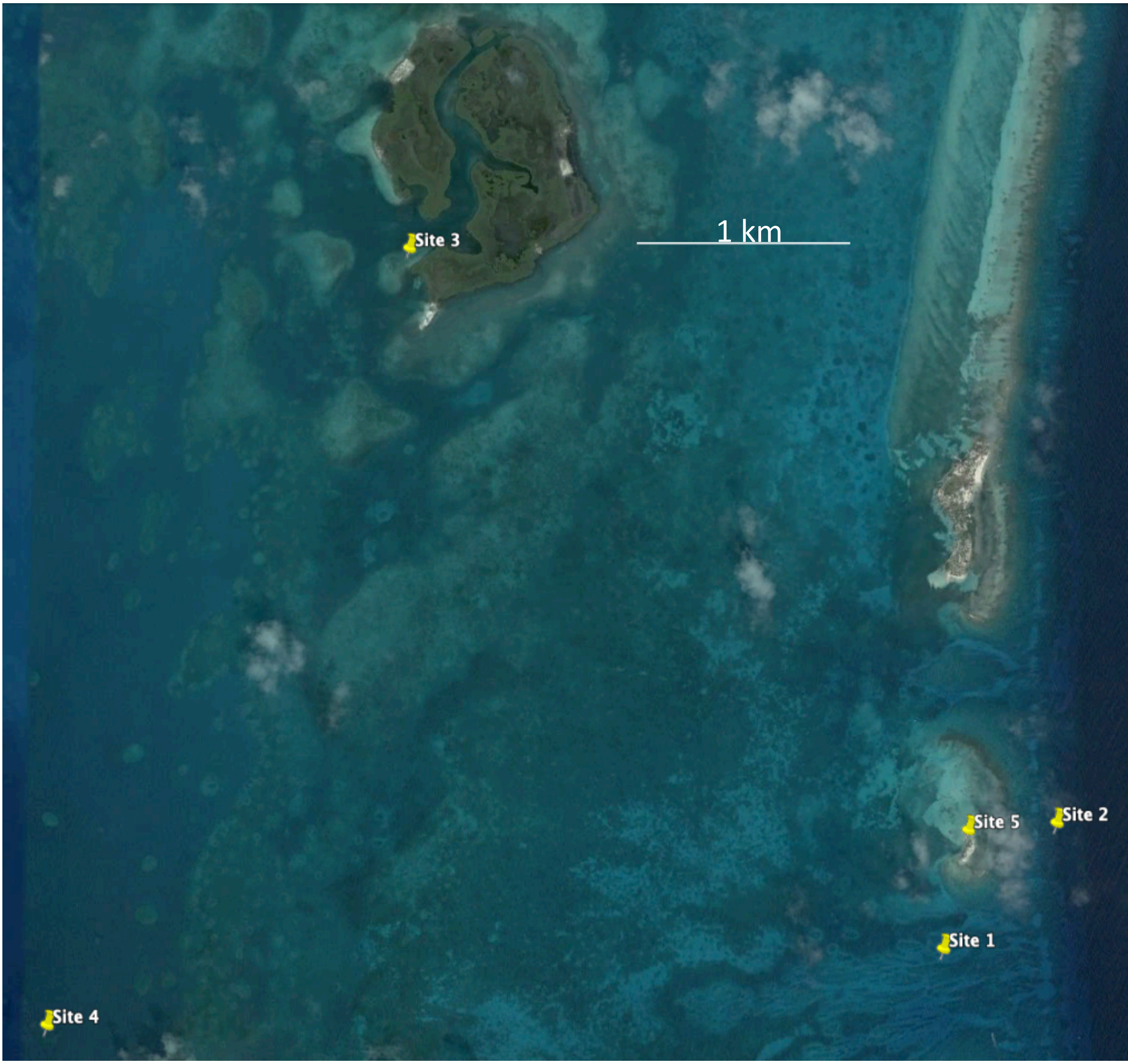


Figure S6. Sediment sample collection site information.

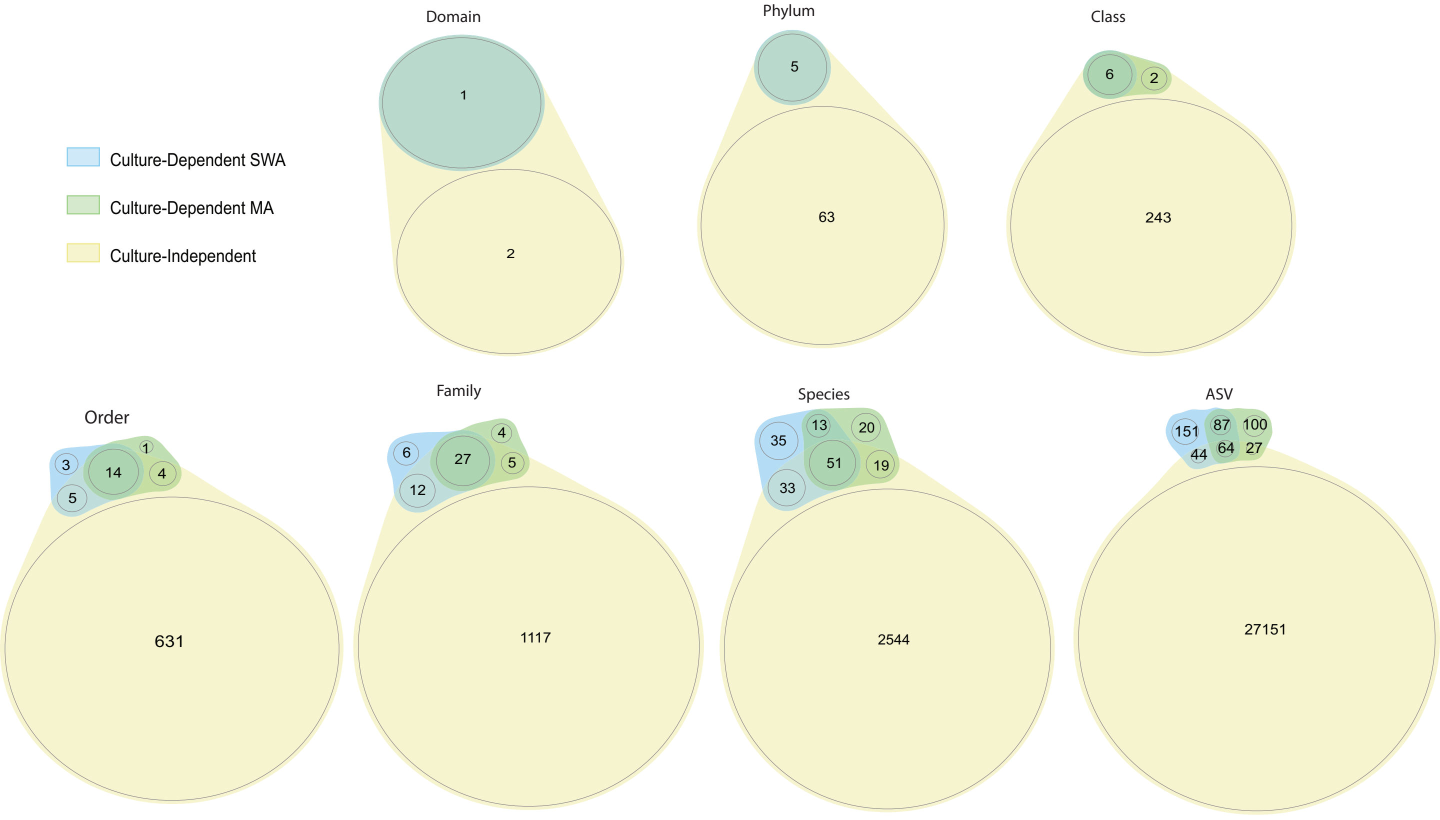


Figure S7. Proportional Venn Diagrams denoting the number of taxa detected with culture-dependent and culture-independent methods (marine agar and seawater agar) across taxonomic levels other than genus.
